# *Spiroplasma* and heat hardening can buffer insect male fertility loss at high temperatures

**DOI:** 10.1101/2025.07.22.666101

**Authors:** Sian Macdonald, Lucy Hayward, Xinyue Gu, Perran A. Ross, Belinda van Heerwaarden

**Author notes:** These authors should be considered joint first authors (Sian Macdonald, Lucy Hayward).

## Abstract

Insects’ upper thermal limits for survival, activity, and fertility have been used to assess vulnerability to climate change, yet heritable endosymbionts - present in over 70% of insect species - are often overlooked. While emerging research suggests some endosymbionts can increase thermal tolerance, their effects on upper lethal and fertility thermal limits has rarely been investigated. Additionally, short-term exposure to sub-lethal high temperatures (heat hardening) can increase insect heat tolerance, but its impact on male fertility is unclear and potential interactions with endosymbionts has not been explored. Here, we investigate whether the endosymbiont *Spiroplasma poulsonii* and heat hardening influence upper survival and fertility thermal limits in a native widespread host of *Spiroplasma* (*Drosophila hydei*) and a novel host (*Drosophila birchii*), a rainforest-restricted species with low heat tolerance. Heat hardening generally improved survival and fertility following a heat shock. The presence of *Spiroplasma* increased survival and fertility of *D. hydei* males following heat-shock, while in *D. birchii,* it did not enhance survival but protected male fertility after heat-shock and modulated hardening responses at sub-lethal temperatures. While protective effects varied across species, sex, and trait, both *Spiroplasma* and hardening significantly buffered male fitness loss during heat shock in both host species and halved heat exposure risk under current and predicted climate change in the rainforest restricted *D. birchii*. These findings highlight that, beyond generating novel phenotypic variation, endosymbionts can interact with plastic responses to heat stress, emphasising the importance of accounting for endosymbiont-mediated effects on thermal tolerance when assessing insect vulnerability to climate change and exploring strategies to manipulate insect thermal tolerance.

## Introduction

Comparisons of species’ upper survival or activity thermal limits (critical thermal maxima: CT_max_) to current habitat temperatures have been utilised to explore vulnerability to climate warming (Diamond et al. 2012, Kellermann et al. 2012, Hoffmann et al. 2013, Pinsky et al. 2019, Clusella-Trullas et al. 2021). However, fertility (particularly male fertility) can be lost at temperatures lower than the lethal threshold for development or adults surviving a heat shock (David et al. 2005, Porcelli et al. 2017, Parratt et al. 2021, van Heerwaarden and Sgrò 2021, Weaving et al. 2024, Pekľanská et al. 2025). In *Drosophila*, these fertility thermal limits have been shown to align more closely with habitat temperatures than CT_max_ (Parratt et al. 2021, van Heerwaarden and Sgrò 2021) and closer to the mean temperature of extinction than other heat tolerance traits such as CT_max_ or survival (van Heerwaarden and Sgrò, 2021). This suggests that fertility loss at high temperatures may play a key role in shaping insect species distributions and determining vulnerability to climate change.

While studies suggest that insects are living in habitats close to their upper critical and fertility thermal limits, they largely ignore phenotypic plasticity and heritable microbes, which might provide a potential buffer or source of additional variation (Wernegreen 2012, Corbin et al. 2017, Dunn 2017, Renoz et al. 2019, Hector et al. 2022). Around 70% of insects are predicted to carry heritable endosymbionts, which are typically passed from mothers to their offspring via the egg (Weinert et al. 2015, Sazama et al. 2019). Some endosymbiont-host relationships have evolved together for millions of years, while others are more recent, with new interactions expected to arise as species move to new areas due to climate change (Wernegreen 2017, Renoz et al. 2019). Recent evidence suggests that endosymbionts can influence how their host responds to thermal stress. Some studies have found endosymbionts can decrease insect resistance to heat stress (Zhu et al. 2021, Gu et al. 2023, Hu et al. 2023), while others have shown that endosymbionts also have the potential to increase host survival after a heat stress (Montllor et al. 2002, Brumin et al. 2011, Gruntenko et al. 2017, Burdina et al. 2021, Majeed et al. 2022). Although there is growing evidence that endosymbionts impact insect heat tolerance, most studies measure heat_-_shock survival, despite emerging evidence suggesting that fertility at high temperatures may be an important predictor of climate change vulnerability (Walsh et al. 2019, Parratt et al. 2021, van Heerwaarden and Sgrò 2021). While one study recently revealed some *Wolbachia* strains can reduce male fertility of their host *Drosophila simulans* under heat stress (Ferguson et al. 2024), it is unclear whether endosymbionts generally impact their host’s fertility at high temperatures.

Although phenotypic plasticity can increase CT_max_ through acclimation (long term exposure) or hardening (short term non-lethal heat shock) (Gunderson and Stillman 2015, Sgrò et al. 2016, van Heerwaarden et al. 2016, Pottier et al. 2022), there is limited data on plasticity for male fertility thermal limits (Jørgensen et al. 2006, Sales et al. 2018, Walsh et al. 2021). One study found that heat hardening at 38 °C for 1 hour significantly decreased the proportion of *D. buzzatii* males becoming sterile following a subsequent heat shock eight hours later (Jørgensen et al. 2006). In contrast, a study on *D. virilis* found that heat hardening had no effect on male fertility on heat shocked pupae or adults, despite enhancing survival (Walsh et al. 2021). However, this study did not include a recovery period, which may be important for inducing hardening responses (Krebs and Loeschcke 1994, Dahlgaard et al. 1998, Kellett et al. 2005, Krebs and Loeschcke 2008), although this has not been examined in relation to fertility. It remains unclear how widespread beneficial hardening responses are, or the extent to which hardening can mitigate male fertility loss under high temperature stress.

While there is evidence that endosymbionts can influence insect plastic responses in life history traits (Da et al. 2016), it is not clear whether endosymbionts can influence plastic responses such as heat hardening in insects. Although Maes et al. (2012) found that both cold acclimation and antibiotic curing of the endosymbionts *Wolbachia* and *Rickettsia* affected the freezing tolerance of *Macrolophus pygmaeus* bugs, the endosymbionts did not appear to influence the bugs’ acclimation capacity. To our knowledge, no studies have examined the effect of endosymbionts on insect heat hardening capacity.

Climate change is predicted to increase the instances of novel species interactions, which could lead to possible host shifts for endosymbionts (Pecl et al. 2017). Interactions between an endosymbiont and a novel host could result in a rapid evolutionary shift within the host species and potentially modify its roles in the ecosystem (Himler et al. 2011). Consequently, understanding whether effects of endosymbionts on native hosts transfer to novel hosts will assist in predictions for species responses to climate change, as well as potentially providing novel management strategies, as endosymbionts can be artificially transferred across hosts though microinjection or sharing the same environments (Huigens et al. 2004, Hoffmann et al. 2011, Brown and Lloyd 2015, Gu et al. 2023). While some effects of endosymbionts on native hosts can be conferred to novel hosts (Tsuchida et al. 2011, Haselkorn and Jaenike 2015), this is not always the case. For example, Sasaki et al. (2005) found a *Wolbachia* strain that induces cytoplasmic incompatibility in its native host, the almond moth (*Cadra cautella*), induces a male killing effect when transferred into a novel host, the Mediterranean flour moth (*Ephestia kuehniella*). It is unknown whether effects on thermal traits can be transferred to novel host species.

In this study, we explored whether the endosymbiont *Spiroplasma poulsonii* influences heat shock survival, fertility and plasticity of its native host, *Drosophila hydei,* and a novel host by transinfection, *D. birchii,* that is restricted to tropical rainforests and is sensitive to heat stress (Schiffer and McEvey 2006, Kellermann et al. 2012, van Heerwaarden and Sgrò 2021). While some *Spiroplasma* strains can induce a male-killing phenotype in *Drosophila*, ladybird beetles, spiders and a butterfly species (Tinsley and Majerus 2006, Haselkorn 2010), the *S. poulsonii* strain found in *D. hydei* does not (Osaka et al. 2008, Haselkorn et al. 2013). Past studies have found no effect of carrying *S. poulsonii* on fecundity and egg-to-adult survival at 25 °C in *D. hydei* (Xie et al. 2011). However, whether *S. poulsonii* influences the thermal tolerance of its host has not been tested. While both plasticity and the presence of endosymbionts can influence insect thermal tolerance, no study has determined their relative effect and whether they interact. Here, we provide the first test of how the endosymbiont *Spiroplasma* affects heat shock survival, fertility, and plasticity in both its native host, *Drosophila hydei*, and a novel, heat-sensitive host, *D. birchii*, to better understand the role of endosymbionts in shaping thermal tolerance, plasticity and climate change resilience.

## Materials and Methods

Detailed methods for fly collection, husbandry, screening and strain identification of *Spiroplasma*, creation of *Spiroplasma* positive and cured lines, and density control prior to experiments are provided in the Supplementary Methods. Briefly, *Drosophila hydei* and *D. birchii* iso-female lines were established from field collections and maintained in discrete generations at 25 °C. The same naturally *Spiroplasma* positive *D. hydei* line was used to establish *Spiroplasma* positive *D. birchii* lines via microinjection. Cured lines were generated using antibiotic treatment and verified by qPCR. Prior to experiments, larval density was controlled and the presence or absence of *Spiroplasma* was confirmed in each line.

### Male and female adult heat-shock lethal thermal limit

Unhardened and heat hardened (see below) adult *D. hydei* and *D. birchii* females and males were exposed to different sublethal and lethal heat shock temperatures to assess the effect of *Spiroplasma* and heat-hardening on their heat-shock survival. *D. hydei* males and females were assessed in separate generations, while *D. birchii* males and females were assayed in the same generation, but different runs on the same day. Flies were separated by sex under a dissecting microscope using CO_2_ and were given at least 48 hours to recover from any adverse effects from the anaesthetic before heat stress (MacMillan et al. 2017). Vials of separated female *D. hydei* were given extra males to continue mating (which were removed without CO_2_ just before the assay), as males become reproductively mature later than females (Markow 1985) (see below).

Heat hardening involved distributing approximately 20 individual flies into 5-10 vials (100_-_200 total) which contained 3mL of fly food media and a piece of card to avoid flies getting stuck in the media under heat. For *D. hydei*, vials were then submerged in a heated water bath at a constant temperature of 37 °C for 1-hour. For *D. birchii*, vials were placed in a controlled temperature incubator (PHCbi) set to 35 °C for 1-hour. Afterwards, vials were placed at 25 °C for 23 hours to recover before the heat-shock. The heat hardening treatment temperatures were selected based on pilot experiments for *D. birchii* and published studies for *D. hydei* (Kellermann and Sgrò 2018, van Heerwaarden et al. 2024), which induced the highest hardening response for adult CT_max_. Survival following all hardening treatments was 100%, to ensure that any effects are the result of plasticity, rather than selection on heat survival.

Heat shock was applied by placing five adult males/females into 0.5mL Eppendorf tubes, which were placed into Biometra Trio 48 thermocyclers (Biometra, Göttingen, Germany) set to different constant temperatures for 1-hour (see below), which were chosen from pilot experiments on each species/sex to include temperatures where survival ranged from 0 – 100%. After exposure to the heat shock, the Eppendorf tubes were opened and placed in a vial with 12mL of food media. The vials were laid on their sides and placed at 25°C. The flies were given 24 hours to recover before they were scored for survival (signs of movement) after heat exposure.

For *D. hydei*, 8-day-old hardened and unhardened males were heat-shocked for 1-hour at 37, 38, 38.5, and 39°C. At 37, 38, and 38.5°C, 20 replicates of 5 flies (100 males per hardening treatment/temperature) were used, while at 39°C, 10 replicates of 5 flies each (50 males per hardening treatment/temperature) were heat-shocked (total n = 1600). Ten-day old hardened and unhardened *D. hydei* females were heat-shocked for 1-hour at 37.5, 38, 38.5 and 39 °C. Fifty females (10 replicates of 5 flies) were stressed per temperature, per treatment. Older females were stressed because we were interested in exploring the fertility of inseminated females (see below), and *D. hydei* males take at least seven days to reach reproductive maturity at 24 °C (Markow 1985).

For *D. birchii*, 7-day-old hardened and unhardened males and females were heat-shocked for 1-hour at 35.5, 36, 36.5, 37, and 37.5°C. Ten replicates of 5 flies (50 flies per sex/hardening treatment) were exposed to each temperature.

### Male adult heat-shock fertility thermal limits

To determine whether *Spiroplasma* or heat-hardening can buffer against heat induced sterility, and whether *Spiroplasma* influenced hardening capacity, male fertility was assessed following exposure to sub-lethal temperatures above. Twenty-four hours after exposure, thirty of the surviving males per line, per hardening treatment and temperature treatment were randomly chosen. Males were distributed into individual vials with 12mL of food media and were given one (*D. birchii*) or two (*D. hydei*) virgin females (from the same batch of density-controlled flies) which had not experienced the heat shock exposure. The fly crosses were then placed at 25 °C and given seven days to mate before both the male and females were removed. After two weeks, the vials were observed for signs of larval activity. If larvae or pupae were present, males were recorded as fertile. Based on effects of *Spiroplasma* and hardening observed in *D. hydei* (which was assessed first), offspring from *D. birchii* were also counted to achieve higher-resolution data.

### Female adult heat-shock stored sperm fertility thermal limit

Previous studies have found that *Drosophila* and flour beetle females do not appear to be sterilised by acute heat shock, but sperm stored within females are sensitive to heat (Sales et al. 2018, Walsh et al. 2022). To explore whether *Spiroplasma* influences stored sperm fertility in females, mated females of each species were observed to see if they were fertile following exposure to heat shock. Twenty-four hours after exposure to sub-lethal/lethal temperatures, 30 of the surviving females from each temperature, line, and hardening treatment were selected to assess whether they were still fertile. Single females were distributed into individual vials with 12mL of food media, placed at 25°C and given five days to lay. After five days, 10 females were randomly selected and transferred to new vials and a new male was added to determine whether the heat shock had sterilised the female, rather than influencing stored sperm viability (although female fertility recovery from heat stress could have also occurred during this period). These females were then removed from the vial five days later. Two weeks after flies were removed, the vials were observed for signs of larvae activity, and if there were larval track marks, larvae or pupae present, the female was scored as fertile for each temperature/time point. Like males, offspring from *D. birchii* were also counted to achieve higher-resolution data.

### Statistical Analysis

To estimate the upper thermal limits of hardened and unhardened *Spiroplasma* (S+) and cured lines (S-), the *drc* package (version 3.0-1) (Ritz et al. 2015) was used in R studio (2025.05.0) to produce three-parameter log-logistic dose-response models. The effective dose function was then used to estimate LT_50_, the temperature at which the survival drops to 50% (Parratt et al. 2021, van Heerwaarden and Sgrò 2021) and compared statistically using the EDcomp function (drc package) (Ritz et al. 2015).

To explore the effect of *Spiroplasma* and heat hardening on fertility (proportion fertile (binomial, logit distribution) and offspring count (*D. birchii* only) (negative binomial or gaussian distribution) following heat shock at each temperature, a general linear model (GLM) using the *lme4* package (version 1.3-959)(Bates et al. 2015) or glmmTMB package (version 1.1)(Brooks et al. 2017) for negative binomial was fitted in R studio. Distributions were selected based on model fit diagnostics using the DHARMa package (version 0.4.6) (Hartig 2024), which indicated they provided the best fit to the data. Independent variables (*Spiroplasma* status and hardening treatment) were designated as fixed effects. Significance of fixed effects was tested using Wald χ^2^ tests (Bolker et al. 2009) in the car package (version 3.0-12) (Fox and Weisberg 2019). To determine whether *Spiroplasma*/ hardening treatments were significantly different at each temperature, we used Tukey’s pairwise comparisons of estimated marginal means (emmeans) using the emmeans package (version 1.8.7) (Lenth 2017). Survival, sterility and fertility plots were produced using the ggplot2 package (version 3.3.5) (Wickham 2016).

As survival and reproduction together determine an organism’s fitness and ability to persist and contribute to future generations, assessing their combined impact will be important for understanding population viability. To evaluate the overall effect of *Spiroplasma* and heat hardening on combined fitness (proportion surviving and fertile/number of offspring), we used a non-parametric bootstrapping approach (in R studio) to assess the uncertainty in the product of survival and fertility (fitness), as not all survivors were assessed for fertility at all temperatures. For each temperature-treatment combination, we performed 10,000 bootstrap resamples. This involved sampling with replacement from the observed data for both survival and fertility and computing the product of these two values for each resample. This generated a distribution of fitness values from which we calculated the mean, as well as the 2.5% and 97.5% percentiles to estimate the 95% confidence interval for each group, which we used to determine whether groups were significantly different from each other (non-overlapping confidence intervals).

### Climate modelling

To explore whether beneficial effects of hardening and *Spiroplasma* on fitness at high temperatures translates to a reduction in overheating risk, we decided to focus on *D. birchii,* as they have a restricted distribution and we have extensive collection data across their distribution in Australia (Schiffer and McEvey 2006). Given that females were poor at storing sperm after heat stress (but heat stressed females were fertile when mated to fertile males) (see results below), we decided to focus on male fitness to explore the effects of *Spiroplasma* and heat hardening on heat exposure risk for *D. birchii*.

We first sourced hourly temperature data for Queensland, Australia (latitude -10° to -25 °S; 140 ° - 145 °E) from the ERA5-Land dataset (2 m above ground, downloaded from the Copernicus Climate Data Store; https://cds.climate.copernicus.eu/) for January 2025 to get an estimate of current overheating risk. To simulate mid-century conditions (2050 - 2070) under the SSP2 4.5 scenario, we added +1°C to each hourly value, reflecting projected global mean temperature increases relative to 2025 (IPCC 2023).

We defined thermal exceedance as the number of hourly timepoints exceeding the critical thermal limit for male composite fitness (survival × fertility), derived from dose–response models (drc package, see above) using average fitness points at each temperature from Figure 3. These thresholds were calculated for each treatment (*birS-* unhardened; *birS-* hardened; *birS+* unhardened; *birS+* hardened) as the effective dose (ED50) at which fitness dropped to 50% of the control (non-heat stressed) (see Figure S1).

To quantify spatial patterns of thermal exceedance, we used the R packages terra (version 1.8-50) (Hijmans 2023) and sf (version 1.0-21) (Pebesma 2018) for raster and spatial vector operations. The Australia shapefile was downloaded using the rnaturalearth package (version 1.01.1) (South 2017) and converted to sf format for plotting. For each treatment, we generated binary hourly rasters (1 = temperature above threshold, 0 = below) and created binary layers indicating whether each pixel exceeded the threshold at least once. These maps were used to calculate the proportion of suitable habitat exposed to exceedance.

Climatically suitable habitat for *D. birchii* was identified using the maximum entropy method (MaxEnt) (Phillips et al. 2006), implemented in the maxnet package (version 0.1.4) (Phillips et al. 2017). Occurrence records were sourced from Schiffer and McEvey (2006). Environmental predictors (BIO1, BIO5, BIO6, BIO12) were sourced from WorldClim v2.1 (Fick and Hijmans 2017) at a 30 arc-second resolution (∼1 km spatial resolution at the equator). To restrict background sampling to ecologically relevant areas, we cropped environmental layers to a 2° buffer around known *D. birchii* occurrence points (VanDerWal et al. 2009) using terra. We extracted predictor values for 10,000 background points, trained the MaxEnt model with default settings using cloglog output, and evaluated performance using AUC using the pROC package (version 1.18.5) (Robin et al. (2011)). Binary suitable/unsuitable maps were generated using the 10th percentile training presence threshold (suitability > 0.26), and cross-validated using minimum training presence as a sensitivity test. To quantify risk, we masked exceedance maps with the binary suitable habitat maps using terra, calculated total exceedance hours, and estimated the percentage of suitable habitat pixels that experienced at least one hour above the threshold.

## Results

### *Spiroplasma* enhances heat hardening responses for heat-shock survival in *D. hydei* males, but not females

Unhardened and hardened (37°C for 1-hour) *D. hydei* males and females, with or without *Spiroplasma* (*hyS*+, *hyS*-), were subjected to a 1-hour heat shock at different temperatures to assess whether prior heat-hardening and *Spiroplasma* influenced male and female upper lethal thermal limits. *Spiroplasma* had little effect on survival in unhardened males; the difference in the LT_50_ values between unhardened *hyS*+ males (LT_50_ = 38.00 °C, 95% CI = 37.97, 39.04) and unhardened *hyS*- males (LT_50_ = 37.89 °C, 95% CI = 37.74, 39.03) was small (Δ = 0.11 °C) and not significant (ED comparison, *t* = -1.56, *P* = 0.12) (Figure 1A). However, in hardened males, *Spiroplasma* had a significant, positive effect on survival (ED comparison, *t* = -7.01, *P* < 0.001), increasing LT_50_ by 0.31 °C (hardened *hyS*+ males LT_50_ = 38.52 °C, 95% CI = 38.48, 38.56; hardened *hyS*- males LT_50_ = 38.21 °C, 95% CI = 38.14, 38.28) (Figure 1A). Heat hardening had a significant, positive effect on male survival in both *hyS*- (ED comparison, *t* = -4.1, *P* = <0.001) and *hyS*+ males (ED comparison, *t* = -18.8, *P* = <0.001). Heat hardening had a stronger positive effect in *hyS*+ males, increasing their LT_50_ by 0.49 °C compared to 0.31 °C in *hyS*- males (Figure 1A).

**Figure 1.**
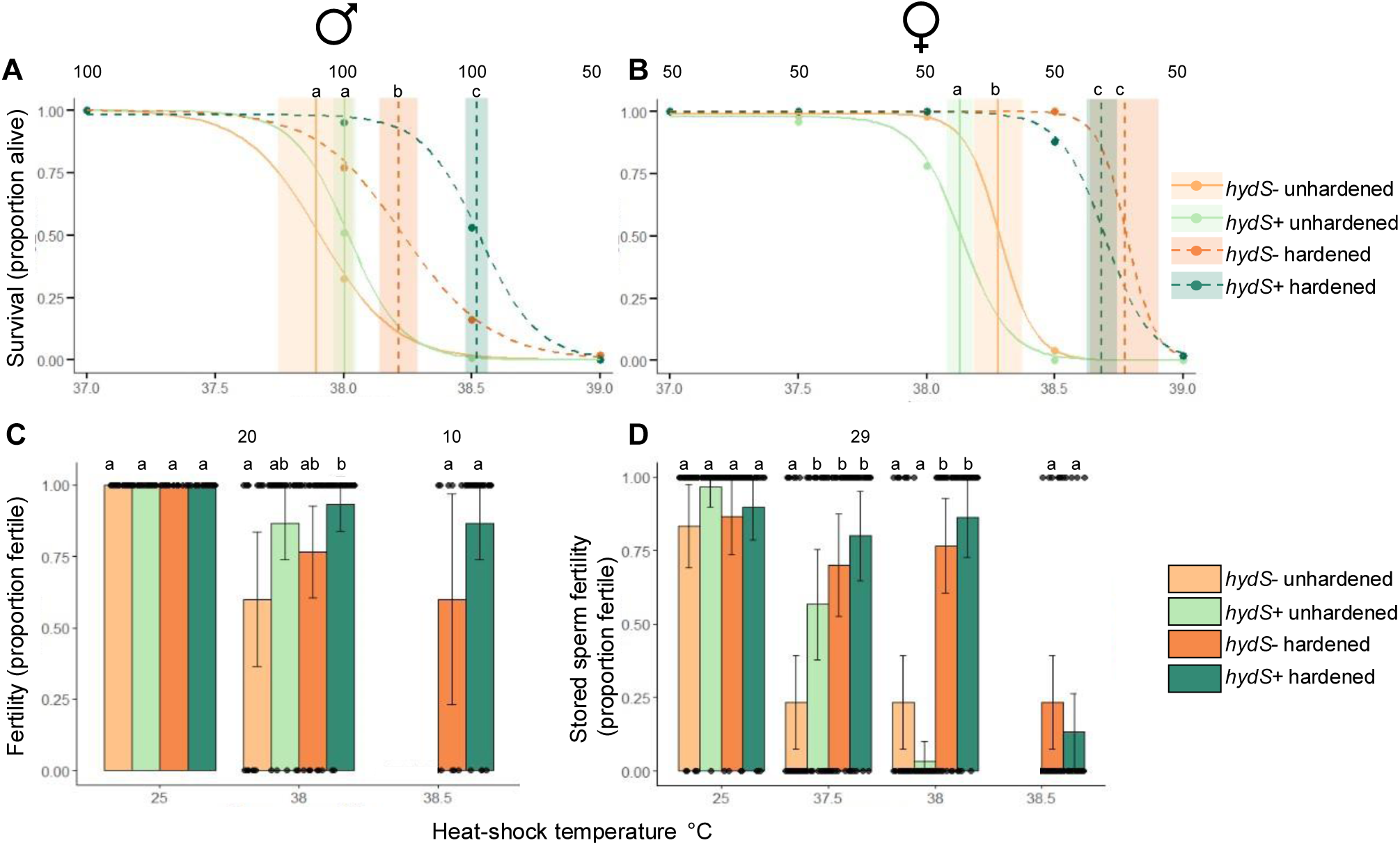
The effect of heat hardening (1 hour at 37 °C) and *Spiroplasma* on *D. hydei* survival (A and B) and fertility (C and D) on flies exposed to 1-hour heat shocks between 37 and 39 °C. Panels A and C show effects on males and panels B and D show effects on mated females. In A and B, circles are the average proportion survival at each temperate (across 10 or 20 replicates with 5 flies per replicate, total N specified above temperature points) and the lines represent the fitted dose response model. Vertical lines represent the LT_50_ for each treatment and the shaded areas are the 95% confidence intervals (CIs). In C – D, bars represent average values, and the capped lines are 95% CIs (across 30 replicates, unless specified above bars). Dots represent data from individuals. Letters above bars indicate treatments that are significantly different from each other (*P* < 0.05, Turkey adjusted).

In contrast to findings in males, LT_50_ was significantly lower (Δ = 0.16 °C, ED comparison, *t* = 3.32, *P* = 0.001) in unhardened *hyS*+ females (LT_50_ = 38.12 °C, 95% CI = 38.07, 38.17) compared with unhardened *hyS*- females (LT_50_ = 38.28 °C, 95% CI = 38.20, 38.35), but there was no significant difference (Δ = 0.09 °C, ED comparison, *t* = 1.45, *P* = 0.150) between hardened *hyS*+ (LT_50_ = 38.68 °C, 95% CI = 38.62, 38.74) and hardened *hyS*- females (LT_50_ = 38.77 °C, 95% CI = 38.66, 38.87) (Figure 1B). Like males, hardening significantly improved the LT_50_ of both *hyS*+ (ED comparison, *t* = 13.91, *P* < 0.001) and *hyS*- females (ED comparison, *t* = 7.59, *P* < 0.001), but responses were similar, increasing the LT_50_ of *hyS*+ females by 0.56 °C and by 0.49 °C in *hyS*- females (Figure 1B).

### *Spiroplasma* and hardening can protect *D. hydei* fertility loss after heat-shock

Surviving *hyS*-/ *hyS*+ and unhardened/hardened males heat shocked at 38 and 38.5 °C and untreated at 25 °C were assessed for fertility following exposure to heat shock. All males were fertile at 25 °C, regardless of hardening treatment (Figure 1C). *Spiroplasma* had a significant, positive effect on the ability of males to maintain fertility at 38 °C (GLM: ꭕ^2^ 1,107 = 8.06, *P* = 0.005), with 90% of *hyS*+ males fertile compared with 70% of *hyS*- males (Figure 1C, Table S1). There was no effect of hardening on the proportion of fertile males at 38 °C (ꭕ^2^ 1,107 = 2.323, *P* = 0.127) and no interaction between *Spiroplasma* and hardening (Figure 1C, Table S1). At 38.5 °C, only hardened individuals survived and could be assessed for fertility (although there were fewer *hyS*- males that were assessed as they died between the 24-hour assessment and fertility matings, so we had less statistical power to detect significant differences). The average proportion of fertile males was higher for *hyS+* males (0.87) compared to *hyS*- males (0.60) (Figure 1C), but this difference was not significant (ꭕ^2^ 1,38 = 3.012, *P* = 0.83) (Table S1).

Surviving *D. hydei* females carrying stored sperm were also assessed for fertility following exposure to heat shock at 37.5, 38 and 38.5 °C (and unstressed controls at 25 °C) to investigate whether *Spiroplasma* and hardening impact female stored sperm fertility. Ninety percent of unhardened females with stored sperm produced offspring at 25 °C, indicating that most females had mated successfully (Figure 1D). There was no effect of *Spiroplasma* (GLM ꭕ^2^ 1,117 _=_ 2.21, *P* = 0.137) or hardening (ꭕ^2^ 1,117 = 0.09, *P* = 0.767) at 25 °C, suggesting that neither influenced the ability to store sperm in females that were not heat_-_shocked (Figure 1D; Table S1).

Both *Spiroplasma* (ꭕ^2^ 1,117 = 6.69, *P* = 0.009) and hardening (ꭕ^2^ 1,117 = 16.28 *P* < 0.001) had a significant, positive effect on the ability of *D. hydei* females to maintain stored sperm fertility when females heat shocked at 37.5 °C, and there was no interaction between hardening and *Spiroplasma* (Figure 1D; Table S1). Unhardened *hyS*- females had significantly lower stored sperm fertility rates (0.23) than unhardened *hyS*+ females (0.57), and hardened *hyS-* females (0.70) (Figure 1D). Although the proportion of fertile hardened *hyS*+ females (0.80) was higher than unhardened *hyS*+ females (0.57), this difference was not significant (Figure 1D, Table S1). There was also a significant effect of hardening on the proportion of fertile *D. hydei* females heat shocked at 38 °C (ꭕ^2^ 1,116 = 64.49, *P* <0.001), and a significant interaction between *Spiroplasma* and hardening (ꭕ^2^ 1,116 = 6.009, *P* <0.014) (Figure 1D; Table S1). Unhardened females had significantly lower stored sperm fertility rates in both *hyS*+ (0.03) and *hyS*- (0.23) groups compared to hardened females (*hyS*+: 0.86, *hyS-*: 0.80), with hardening improving fertility more in *hyS*+ females (a 0.83 increase) than in *hyS*- females (a 0.54 increase) (Figure 1D). At 38.5°C, only hardened females survived, and fertility rates were low in both *hyS*+ (0.12 and *hyS*- (0.23) females, with no significant difference between them (ꭕ^2^ 1,58 = 1.012, *P* = 0.314) (Figure 1D; Table S1).

### *Spiroplasma* does not improve male heat shock survival in a novel heat sensitive host, but has a positive impact on male fertility at sub lethal temperatures and interacts with heat hardening

Unhardened and hardened (35 °C for 1-hour) *D. birchii* males and females, with and without *Spiroplasma* microinjected from *D. hydei*, were subjected to a 1-hour heat shock at different temperatures (35.5 to 37.5 °C) to assess whether prior heat-hardening and *Spiroplasma* influenced male and female upper lethal and fertility thermal limits in a *Spiroplasma* naïve species. Similar to *D. hydei*, hardening had a significant, positive effect on *D. birchii* male heat-shock survival, increasing the LT_50_ of *birS*+ males by 0.30 °C (ED comparison, *t* = - 6.70, *P* <0.001) and 0.18 °C (ED comparison, *t* = - 2.63, *P* <0.001) in *birS*- males (Figure 2A). In contrast to findings in *D. hydei,* unhardened *birS*- males had a significantly higher LT_50_ (LT_50_ = 36.58 °C, 95% CI 36.52, 36.65) than unhardened *birS*+ males (LT_50_ = 36.47 °C, 95% CI 36.42, 36.52) (ED comparison, *t* = 2.63, *P* = 0.009), improving LT_50_ by 0.11 °C. However, there was no significant difference between hardened *birS*+ males (LT_50_ = 36.77 °C, 95% CI = 36.70, 36.84) and hardened *birS*- males (LT_50_ = 36.76 °C, 95% CI = 36.64, 36.87) (Δ = 0.01 °C; ED comparison, *t* = -0.21, *P* = 0.84) (Figure 2A).

**Figure 2.**
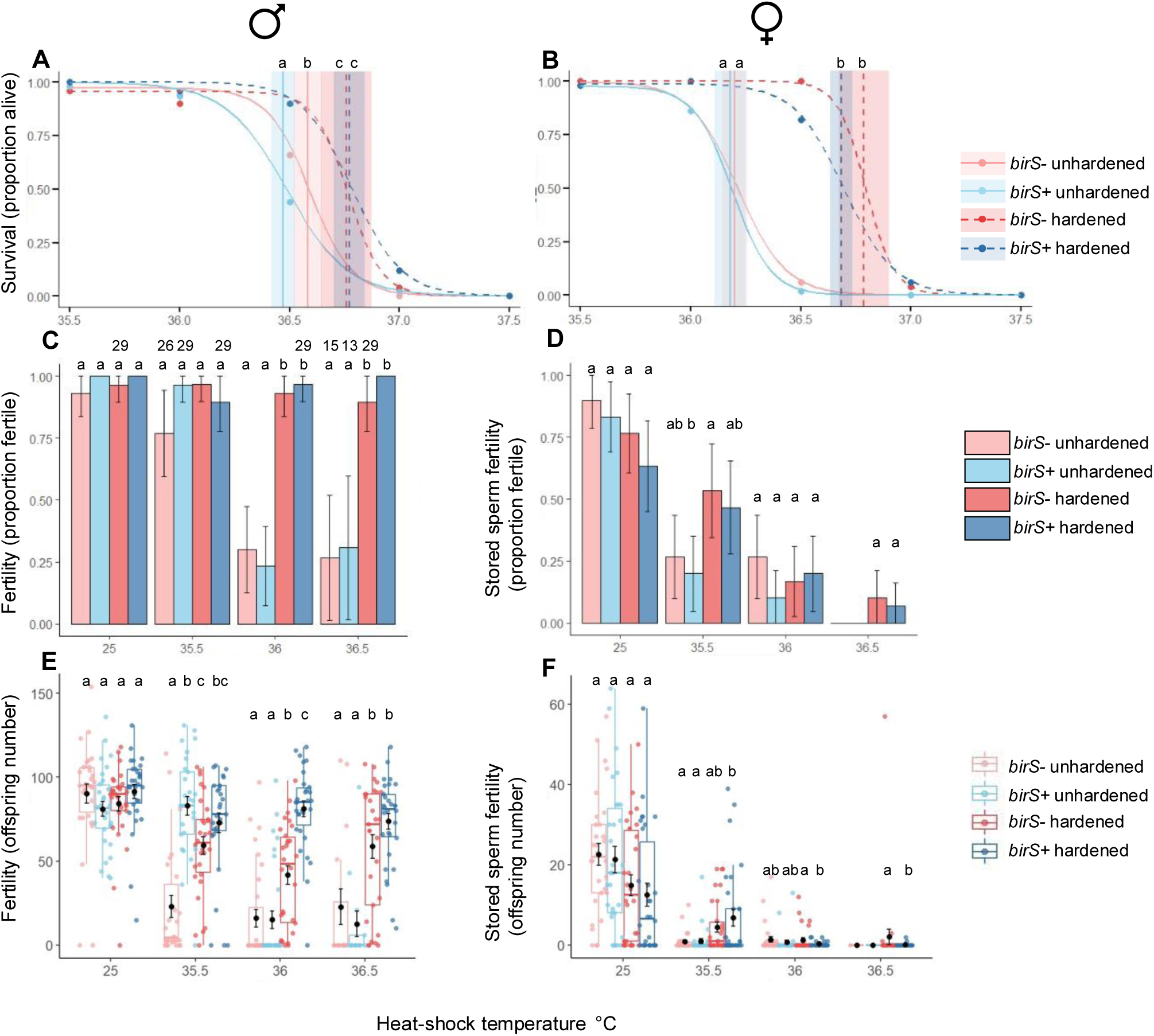
The effect of heat hardening (1 hour at 35 °C) and *Spiroplasma* on *D. birchii* (A and B) survival; (C and D) proportion fertile; and (E and F) fertility (total offspring count of fertile/sterile males/females ) on adult flies exposed to 1-hour heat shocks between 35.5 and 37 °C. Panels A, C and E show effects on males and panels B, D and F show effects on mated females. In A and B, circles are the average proportion survival at each temperate (10 replicates, with 5 flies per replicate) and the lines represent the fitted dose response model. Vertical lines represent are the LT50 for each treatment and the shaded lines are the 95% confidence intervals (CIs). In C – D, bars represent average values, and the capped lines are 95% Cis (across 30 replicates, unless specified above bars). In E and F, the box represents the interquartile range (IQR; 25th to 75th percentile), with the horizontal line indicating the median, whiskers extend to the smallest and largest values within 1.5 × IQR from the lower and upper quartiles, respectively. Jitter points are individual data points. Black dots represent group means, and error bars indicate ±1 standard error of the mean. Letters above indicate treatments that are significantly different from each other (*P* < 0.05, Turkey adjusted).

Hardening also had a significant, positive effect on *D. birchii* female heat-shock survival, increasing the LT_50_ of *birS*+ females by 0.50 °C (ED comparison, *t* = -11.3, *P* < 0.001) and 0.59 °C (ED comparison, *t* = -9.25, *P* = 0.009) in *birS*- females (Figure 2B). *Spiroplasma* had no impact on the LT_50_ of unhardened (*birS*- LT_50_ = 36.20 °C, 95% CI = 36.14, 36.26; *birS*+ LT_50_ = 36.18 °C, 95% CI = 36.11, 36.25, *t* = 0.47, *P* = 0.64) or hardened females (*birS*- LT_50_ = 36.79 °C, 95% CI = 36.67, 36.90; *birS*+ LT_50_ = 36.68 °C, 95% CI = 36.63, 36.74; ED comparison, *t* = 1.63, *P* = 0.10) (Figure 2B).

Surviving unhardened/hardened, *birS*- / *birS*+ males heat shocked at 35.5, 36 and 36.5 °C, and untreated at 25 °C were assessed for fertility following exposure to heat shock. Neither hardening, nor *Spiroplasma* had a significant impact on the proportion of fertile males or offspring number in males that were not heat shocked (control 25 °C), which remained high (>90% fertile, average 85 offspring) across all hardening and *Spiroplasma* combinations (Figure 2C; Table 2). There was a significant effect of hardening on male sterility (proportion of fertile males) at the higher heat shock temperature (36 °C: ꭕ^2^ 1,116 = 67.614, *P* < 0.001; 36.5 °C ꭕ^2^ 1,83 = 45.373, *P* < 0.001), where the proportion of fertile unhardened males declined to around 25%, but remained high (>90%) when males were hardened (Figure 2C; Table S2). There was not a significant effect of *Spiroplasma* on the proportion of fertile males at any heat shock temperature, but there was as significant interaction between *Spiroplasma* and hardening at 35.5 °C (ꭕ^2^ 1,110 = 5.202, *P* = 0.023), with *birS*-, unhardened males showing a lower (but not significantly) proportion of fertile males than *birS+* unhardened males (Figure 2C; Table S2).

Given that some males may be fertile but produce few offspring, we also considered offspring counts (fertility) overall and for fertile *D. birchii* males only. Hardening significantly increased offspring number at all heat-shock temperatures 35.5, 36, and 36.5 °C) when both fertile and sterile males were included in the analysis (Figure 2E, Table S2). In contrast, when only fertile males were considered, hardening significantly increased offspring number at 35.5 °C only (Figure S2, Table S4). This indicates that the strong positive effect of hardening on overall offspring production at higher temperatures is primarily driven by an increase in the proportion of fertile males rather than an increase in productivity among already fertile individuals.

We also found that *Spiroplasma* had a significant positive impact on male fertility at 35.5 and interacted with hardening at 35.5 and 36 °C (Figure 2E, Table S2). At 35.5 °C, unhardened, *birS*- males had significantly lower offspring number than hardened *birS*- males, while offspring number remained high in both unhardened and hardened *birS*+ males for all males and only fertile males (Figure 2E). This pattern was consistent when considering all males (Figure 2E) and when restricting the analysis to fertile males only (Figure S2; Table S4). At 36 °C, when considering both fertile and sterile males, offspring numbers were similar in unhardened *birS*+ and *birS*- males, which were both significantly lower than hardened males (Figure 2E). However, *birS*+ males benefited more from hardening, producing significantly more offspring than *birS*- hardened males (Figure 2E). There was no effect of *Spiroplasma* at 36.5 °C (Figure 2E, S2, Table S2,S4) when considering both fertile and sterile males.

Surviving females carrying sperm were also assessed for fertility following exposure to heat shock at 35.5, 36 and 36.5 °C (and non-heat shocked, control 25 °C) to assess whether *Spiroplasma* and hardening impact female stored sperm fertility (proportion of fertile females/ number of offspring). Eighty-six percent of unhardened females with stored sperm produced offspring at 25 °C, indicating that most females had mated successfully (Figure 2D). There was no effect of *Spiroplasma* on the proportion of fertile females (ꭕ^2^ 1,116 = 1.655, P = 0.198) or offspring number (ꭕ2 1,116 = 0.381, P = 0.537) at 25 °C, suggesting that Spiroplasma had no effect on the ability of non-heat shocked females to store sperm (Figure 2D,F; Table S3). However, hardening had a significant negative impact on both the proportion of fertile females (ꭕ^2^ 1,116 = 4.893, *P* = 0.027) and offspring number (ꭕ^2^ 1,116 = 8.095, *P* = 0.004) at 25 °C (Figure 2D,F; Table S3), indicating that heat hardening had a negative effect on female stored sperm fertility in the absence of a secondary heat stress. At 35.5 °C, only hardening had a significant impact on the proportion of fertile females at (ꭕ^2^ 1,116 =9.413, *P* = 0.002), significantly improving fertility in hardened females in both *birS*+ and *birS*- females from around 20% to 50% (Figure 2D; Table S3). Although there was also a significant beneficial effect of hardening on the overall number of offspring at 35.5 °C (ꭕ^2^ 1,115 = 15.342, *P* < 0.001), hardened females only produced an average of five offspring compared to one for unhardened males (Figure 2F). When considering fertile females only, hardening also had a significant effect, especially in *birS*+ females, which produced on average 15 offspring compared to 3 and 5 offspring in fertile unhardened *birS*- and *birS*+ respectively (Table S4, Figure S2). The proportion of fertile females was low at 36 °C (0.18 average) and 36.5 °C (0.08 average), and neither hardening nor *Spiroplasma* had any significant impact on this or offspring number (Figure 2D, 2F; Table S3). At 36 and 36.5, offspring production in fertile females only was higher in hardened, *birS*- females (Figure S2, Table S4), but this is based on low numbers (N = 5 and N= 3).

### Remating suggests heat shock impairs stored sperm rather than female reproductive function

After six days, a subset of *D. birchii* and *D. hydei* females were remated to non-heat shocked males to assess whether heat shock had directly impacted female fertility or was due to impacts on stored sperm (albeit females may have also recovered fertility during this time). The proportion of fertile female and offspring number (*D. birchii* only) were high across all treatments, suggesting that reductions in fertility were due to impact of temperature on stored sperm (Figure S3). There was no effect of hardening or *Spiroplasma* and no interactions at any temperature on either proportion fertile or offspring number for either species (Figure S3; Table S3). In *D. birchii*, offspring number was much higher in the control 25 °C treatment in remated females with access to a male (average 63.9 offspring: Figure S3C) than the females with stored sperm (average 17.8 offspring; Figure 2F), suggesting that *D. birchii* either reach peak laying after 12 days of age or they are not very good at storing sperm, even when not heat stressed.

### *Spiroplasma* and heat hardening increase composite fitness (proportion survived *and* fertile) following heat shock and increase habitat suitability for rainforest restricted heat sensitive species *D. birchii*

Since *Spiroplasma* and heat hardening affected survival and fertility in varying ways (sometimes both traits, sometimes only one, and occasionally in opposite directions) and both traits combined are essential for fitness, we were interested in assessing their impact on survival and fertility combined. For *D. hydei* males exposed to heat shock at 38 °C, both hardening and *Spiroplasma* improved composite fitness (Figure 3A). Hardening led to a similar improvement in composite fitness in both *hyS*− (Δ = 0.41) and *hyS*+ (Δ = 0.45) males, though unhardened and hardened *hyS*+ males had significantly higher overall fitness than *hyS*- males (unhardened : 0.44 vs 0.18; hardened: 0.89 vs 0.59). At 38.5 °C, hardened males had significantly higher fitness than unhardened males, regardless of *Spiroplasma* status, and hardening improved composite fitness from 0 to 0.10 in *hyS*- males and from 0 to 0.46 in *hyS*+ males, partially rescuing fitness at this highly stressful temperature (Figure 3A). In *D. hydei* females, we also found strong beneficial effects of hardening on composite fitness at all temperatures, but the magnitude and pattern of the response varying depended on temperature and *Spiroplasma* status (Figure 3B). At 37.5 °C, hardening significantly improved composite fitness from 0.23 to 0.70 in *hyS*- females and from 0.54 to 0.80 in *hyS*+ females. At 38 °C, hardening increased composite fitness from 0.2 to 0.77 in *hyS*- females and from 0.03 to 0.87 in *hyS*+ females. At 38.5 °C, where baseline fitness was zero in unhardened *hyS*- and *hyS*+ females, hardening rescued fitness to measurable levels in both groups, with fitness rising to 0.23 in *hyS*- and 0.12 in *hyS*+ females, which were not significantly different from one another.

**Figure 3.**
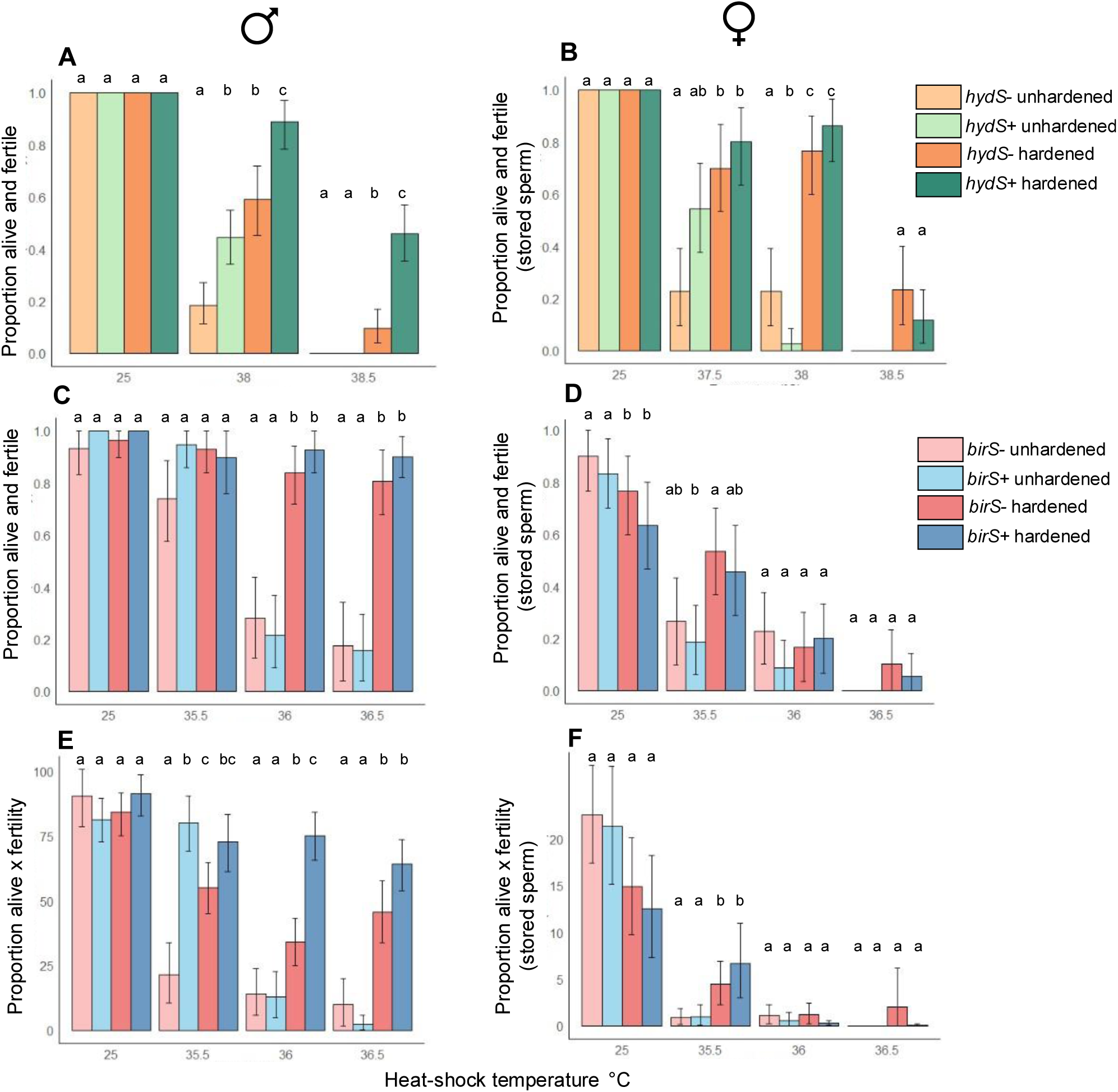
The effect of heat hardening (*D. hydei*: 1 hour at 37 °C; *D. birchii*: 1 hour at 35 °C) and *Spiroplasma* on *D. hydei* (A and B) and *D. birchii* (C, D, E and E) on combined survival and fertility measures in adult flies exposed to 1-hour heat shocks and no heat shock (25 °C). Panels A – D show effects on the proportion that survive and are fertile, while panels E and F show effects on that proportion that survive x offspring number. Panels A, C and E show effects on males and panels B, D and F show effects on mated females. Bars represent average values, and the capped lines are 95% CIs from bootstrap analysis. Letters above indicate treatments that are significantly different from each other (*P* < 0.05, 95% CIs non overlapping).

Consistent with the survival and fertility results considered individually, hardening had a strong beneficial effect on fitness (proportion surviving and fertile/ proportion surviving x offspring number) after heat shock in *D. birchii* males (Figure 3C, E) and small beneficial effects on females (Figure 3D, F). Similar to the fertility results, *Spiroplasma* had a positive effect on male fitness based on offspring output (proportion surviving x offspring number) and interacted with heat hardening (Figure 3E), despite the survival thermal limit being significantly lower in unhardened *birS+* males compared to unhardened *birS–* males (Figure 2A).

When we used these fitness thresholds for *D. birchii* males to explore how *Spiroplasma* and hardening influence heat exposure risk under current summer temperatures and under 1 °C of climate change, we found a marked increase in areas not exceeding upper thermal fitness thresholds across north-east Queenland (Figure 4). These effects translated into increased thermally suitable habitat across *D. birchii’s* predicted geographical range, under both current and future climate scenarios (Figure 5). Under current climate conditions, 34% of suitable habitat exceed male fitness thresholds of unhardened, *birS*- males for at least one hour (Figure 5A). This increased to 55% under 1 degree of climate change (Figure 5E). Hardening and *Spiroplasma* decreased the percentage of suitable habitat exceeding male thermal fitness threshold under current climates to 23% and 27% respectively (Figure 5B, 5C). The combination of hardening and *Spiroplasma* had the biggest impact, decreasing the percentage of suitable habitat exceeding the male fitness thermal threshold to only 15% (Figure 5D). Under a +1 °C warming scenario, hardening and *Spiroplasma* decreased the percentage exceeding male thermal fitness threshold under current climates to 41% and 44% respectively (Figure 5B, 5C). The combination of both hardening and *Spiroplasma* together returned the viable habitat area to similar levels (30%) (Figure 5I) as untreated males under current climates (34%) (Figure 5A).

**Figure 4.**
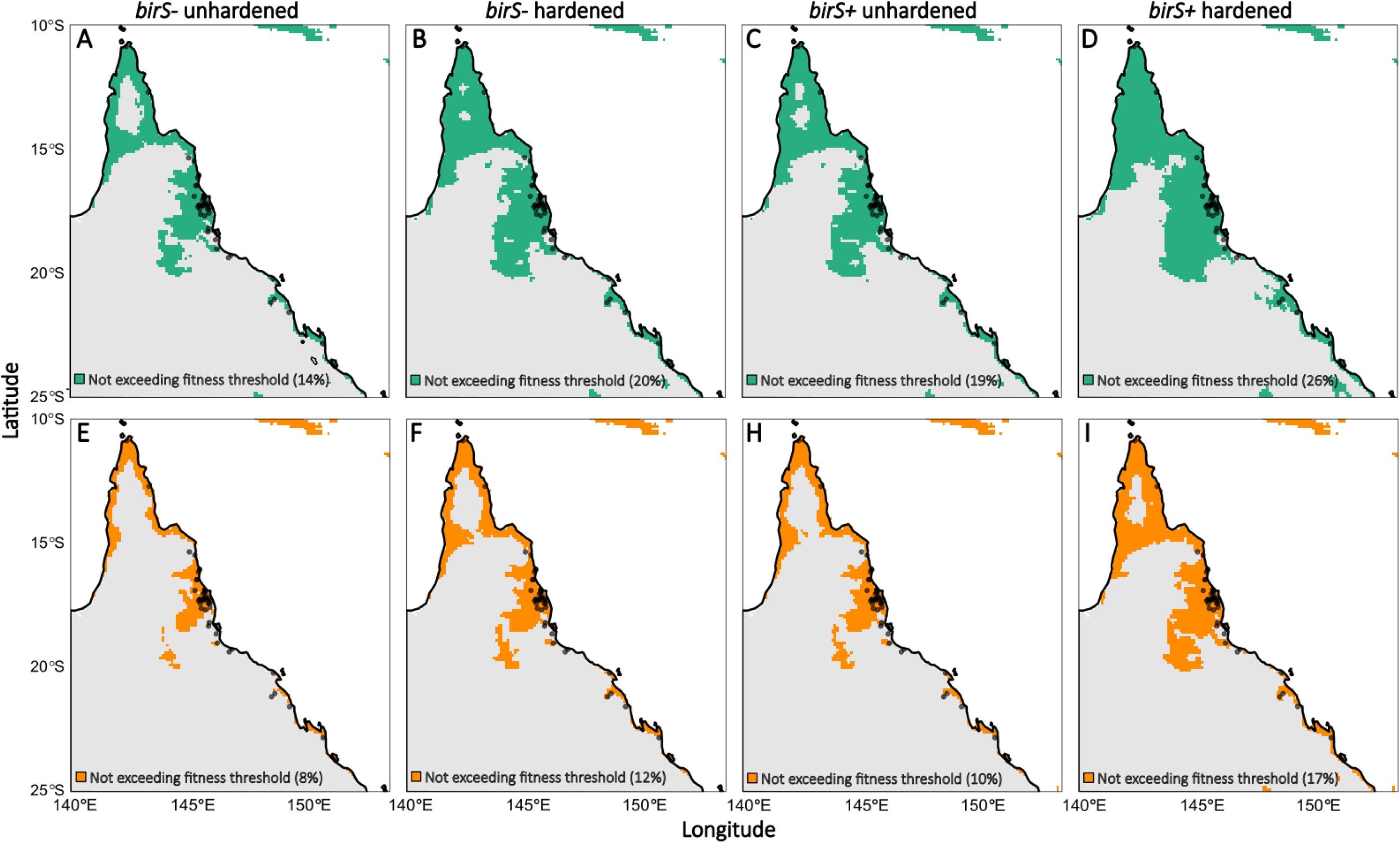
Areas across north-east Queensland that exceed (grey) and do not exceed (green, orange) upper thermal thresholds for fitness (see Figure S1) for at least one hour in the warmest month (January) under current temperatures (A - D, green) and with 1 °C of warming (E - H, orange) for unhardened and hardened *D. birchii* with (*birS*+) and without (*birS*-) *Spiroplasma.* Black dots are known collection locations for *D. birchii* in Australia.

**Figure 5.**
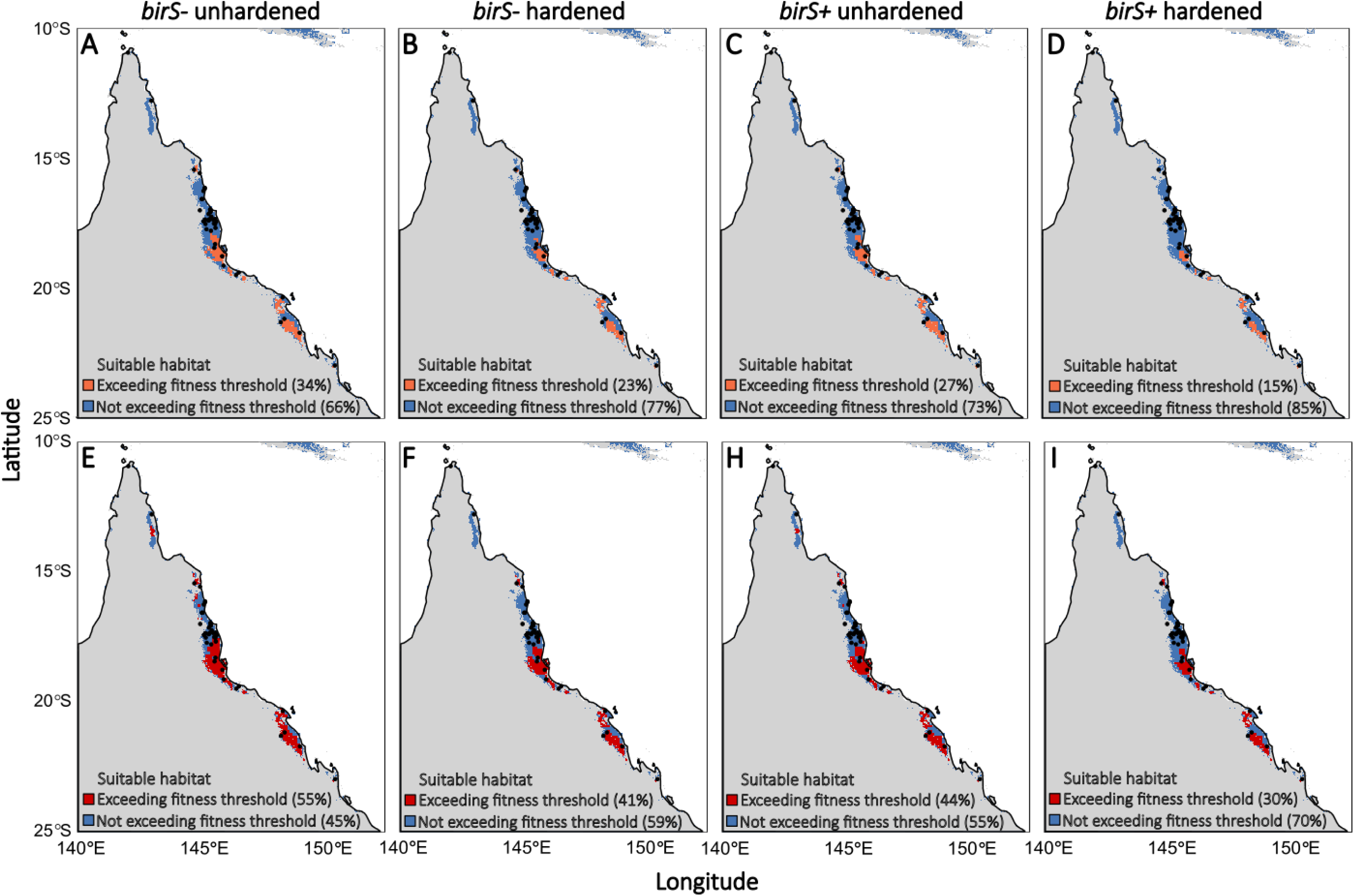
Areas within suitable habitat for *D. birchii* that do not exceed (blue) and do exceed (orange, red) upper thermal thresholds for fitness (see Figure S1) for at least one hour in the warmest month (January) under current temperatures (A–D, orange) and with 1 °C of warning E–H, red) for unhardened and hardened *D. birchii* with (birS⁺) and without (birS⁻) *Spiroplasma*. Grey areas are climatically unsuitable for *D. birchii* based on Maxent modelling (Figure S4). Black dots are known collection locations for *D. birchii* in Australia.

## Discussion

Understanding what drives variation in insect survival and fertility at elevated temperatures is critical for predicting species responses to climate change. Given that evolution and plasticity alone may not be sufficient to buffer species (Gunderson and Stillman 2015, Sgrò et al. 2016, van Heerwaarden and Sgrò 2021, Hoffmann et al. 2023), identifying additional sources of phenotypic variation and mechanisms that shape plasticity may change the way we think about how species respond and adapt to environmental change. We found strong beneficial effects of hardening on survival, fertility and overall fitness after heat shock. We also found that *Spiroplasma* can enhance male heat tolerance, fertility and fitness, and influence plastic responses, revealing a role for endosymbionts as an additional source of phenotypic variation and a potential driver of phenotypic plasticity in thermal tolerance traits. However, when *Spiroplasma* was transferred to a novel, heat sensitive host (*D. birchii*), its effects were more limited, only improving male fertility at high temperatures. Nonetheless, the increased fertility in *D. birchii* males led to a greater number of surviving, fertile males, suggesting that *Spiroplasma* can confer fitness benefits at after heat stress even in new host backgrounds. This resulted in reduced heating risk under current and predicted climate change. Together, our findings highlight the complex and context-dependent ways that plasticity and endosymbionts can interact to influence thermal fitness under climate stress.

### Heat hardening enhances survival and fertility

Plasticity has often been proposed as an important mechanism to buffer climate change in the short-term (Sgrò et al. 2016, Fox et al. 2019), but plastic responses are often small (especially in insects) (Gunderson and Stillman 2015, van Heerwaarden et al. 2016, Weaving et al. 2022). We found substantial hardening responses for survival LT_50_ in both species. In females, hardening improved LT_50_ by 0.52 and 0.54 °C in *Spiroplasma* free *D. hydei* and *D. birchii* respectively. This is more than double the maximum improvement that similar hardening treatments have found for knockdown CT_max_ in females (0.1 °C in *D. hydei*; 0.17 °C in *D. birchii*) (Kellermann and Sgrò 2018). Because most studies exploring the effects of heat hardening or acclimation on survival use a single heat shock temperature, and do not estimate LT_50_, it is unclear whether lethal limits generally have larger plasticity than knock down limits (Gunderson and Stillman 2015, Weaving et al. 2022).

Importantly, hardening also buffered fertility loss, particularly in *D. birchii* males, where it had large impacts on overall fitness. In *D. hydei* males, we did not detect a significant effect of hardening on fertility, potentially due to our use of binary fertility scores (e.g. Walsh et al. 2019, Parratt et al. 2021, Bretman et al. 2024), which may be less sensitive than offspring counts if some heat stressed individuals are fertile, but produce few offspring. While hardening in *D. birchii* males only modestly increased the proportion of fertile individuals at 35.5 °C, it improved offspring production, especially when combined with the presence of *Spiroplasma.* Our findings demonstrating that hardening buffered fertility loss after heat- shock contrast with results on *D. virilis,* where hardening improved survival, but did not protect fertility (Walsh et al. 2021). But our findings align with Jørgensen et al. (2006), who found a strong protective effect of hardening on fertility loss in *D. buzattii*. A key difference may be the inclusion of a recovery period in the current study and Jørgensen’s study; since candidate heat gene expression can peak several hours after stress (e.g. Telonis-Scott et al. 2021), recovery time may be critical to enabling protective responses (Krebs and Loeschcke 1994, Dahlgaard et al. 1998, Kellett et al. 2005, Krebs and Loeschcke 2008). These findings suggest that future studies should assess the effects of recovery periods and consider counting offspring when evaluating male fertility responses.

We also revealed, for the first time, that heat hardening can protect stored sperm fertility after heat shock in females. This is significant, because if females can successfully store sperm and remain fertile after heat-shock, they may partially buffer the impacts of heat stress on male fertility. Consistent with Walsh et al. (2022), who showed that sperm stored in *D. virilis* females was highly susceptible to thermal stress, we found the proportion of unhardened fertile *D. hydei* females carrying stored sperm declined to ∼25% after heat_-_shock at 37.5 and 38 °C in. However, heat hardening increased stored sperm fertility to ∼75%, indicating strong protective effects. In *D. birchii*, hardening also increased stored sperm fertility, though the benefits of hardening were much smaller and only really evident at the mildest heat shock temperature. Interestingly, hardening reduced stored sperm fertility in *D. birchii* females that did not receive a heat shock, suggesting potential fitness costs of the treatment itself (Krebs and Loeschcke 2008). These costs appeared to be short-lived and probably due to heat damage to stored sperm, as we detected no fertility costs one week after heat shock in females allowed to remate. While it is unclear whether limited sperm storage capacity in *D. birchii* (Saxon et al. 2018), underpins these patterns, our results demonstrate that heat hardening can provide a valuable buffer against fertility loss by protecting stored sperm after thermal stress, although the effectiveness of hardening and potential costs may vary across species. Exploring this further will be critical for predicting which species can leverage plasticity to maintain fertility under climate change.

### *Spiroplasma* interacts with heat hardening to enhance male survival and fertility

Both heat hardening and the presence of endosymbionts have been shown to influence thermal tolerance in insects (Montllor et al. 2002, Brumin et al. 2011, van Heerwaarden et al. 2016, Kellermann and Sgrò 2018, Sørensen et al. 2019, Burdina et al. 2021, Majeed et al. 2022), but their relative contributions and whether endosymbionts modulate plastic responses is unclear. Our results demonstrate that heat hardening generally had a greater effect on survival than *Spiroplasma* in both *D. hydei* and *D. birchii*, highlighting the stronger protective role of hardening-induced plasticity in shifting upper thermal limits. In contrast, the relative effects of hardening and *Spiroplasma* on fertility were more variable. In *D. hydei* males, *Spiroplasma* had a greater positive impact on fertility than hardening, whereas in *D. birchii* males and in females of both species (as measured by stored sperm fertility), heat hardening played a more substantial role. Importantly, our findings also suggest that *Spiroplasma* can influence plastic responses. *Drosophila hydei* males with *Spiroplasma* showed enhanced heat survival following hardening, indicating a greater capacity for plasticity compared to males lacking *Spiroplasma*. Similarly, in *D. birchii* males, the effects of heat hardening on fertility were larger in flies carrying *Spiroplasma*, suggesting an interaction between *Spiroplasma* and plasticity in thermal fertility traits.

### Effects of *Spiroplasma* are trait specific in a novel host

Although *Spiroplasma* improved upper thermal survival limits in *D. hydei* males, especially in hardened flies, it had small negative effects on survival in *D. hydei* females and survival in both sexes of *D. birchii*. In contrast, we found consistent positive effects of *Spiroplasma* on male fertility after heat shock in both species. In line with previous studies (Olsen et al. 2001, Anbutsu et al. 2008), our results highlight that endosymbiont effects can depend on host genetic background, likely reflecting a history of co-evolution. Host-dependence complicates general predictions across species and has important implications for novel endosymbiont transfers. As species shift their distributions with climate change, new host– symbiont associations are likely to emerge (Pecl et al. 2017), potentially leading to unpredictable effects on thermal tolerance. From an applied perspective, where endosymbionts are being explored for pest management or climate resilience (Walker et al. 2011, Nazni et al. 2019, Gu et al. 2023), understanding host-specific outcomes is crucial.

### *Spiroplasma* and heat hardening increase composite fitness thermal limit and habitat suitability for rainforest restricted heat sensitive species *D. birchii*

The beneficial effects of heat hardening and *Spiroplasma* were more apparent when we considered their effects on overall fitness (proportion surviving and fertile) following heat shock. Heat hardening and *Spiroplasma* enabled *D. hydei* males to maintain fitness (0.46) at 38.5 °C, a temperature at which unhardened males had zero fitness due to complete mortality. At 38 °C, heat hardening and *Spiroplasma* increased male fitness from 0.18 in unhardened males without *Spiroplasma,* to 0.89 in hardened males carrying *Spiroplasma*. In *D. birchii* males heat hardening was also particularly effective, allowing *Spiroplasma* males to maintain fitness levels comparable to non-heat shocked males right up to their lethal limit. Together, these findings demonstrate that hardening and *Spiroplasma* can dramatically enhance fertility and ultimately fitness at temperatures near or beyond survival limits

Our findings show that both *Spiroplasma* and heat hardening substantially increase the composite fitness thermal limit for the rainforest-restricted, heat-sensitive species *D. birchii*. By increasing the thermal threshold at which male survival and maintain fertility following heat shock, hardening and *Spiroplasma* approximately halved the percentage of predictable suitable habitat exceeding their fitness limits under both current and projected future climates. Although our fitness estimate did not consider accumulated heat damage across multiple heat stress events (Jørgensen et al. 2021, Ørsted et al. 2024), or different hardening treatments (van Heerwaarden et al. 2024), our study demonstrates the substantial potential for symbiont-mediated/plastic responses to buffer fitness loss and expand thermally viable habitat for vulnerable tropical ectotherms.

Although determining the underlying mechanisms is beyond the scope of this study, *Spiroplasma* may influence host thermal tolerance via heat shock proteins (HSPs), which assist in protein refolding and prevent aggregation under heat stress (Feder and Hofmann 1999, King and MacRae 2015). In spider mites, both *Wolbachia* and *Spiroplasma* have been shown to alter HSP gene expression in response to temperature shifts (Zhu et al. 2021), and in aphids, *Spiroplasma* has been shown to regulate HSPs depending on diet (Guidolin et al. 2018). Future work should explore whether similar mechanisms underlie *p oplasma’s* effects on heat tolerance in *D. hydei* and *D. birchii*. Other endosymbiont_-_mediated protective mechanisms include the overexpression of host cytoskeleton genes, as observed with *Rickettsia* in the whitefly *Bemisia tabaci* (Brumin et al. 2011). In *Drosophila*, endosymbionts have also been linked to shifts in metabolite levels associated with thermal stress resistance (Gruntenko et al. 2017, Burdina et al. 2021). Given that *Spiroplasma* can influence host lipid metabolism (Herren et al. 2014), it is also possible that changes in membrane composition contribute to enhanced thermotolerance. Membrane lipids can play a key role in thermal injury and recovery, and their stabilisation is a known component of heat hardening Jóźwiak and and Leyko 1 2), suggesting another potential mechanism by which *Spiroplasma* could modify host heat tolerance via effects on membrane fluidity or integrity.

### Conclusions

Our findings show that *Spiroplasma* can enhance survival and fertility following heat stress, although protective effects varied depending on species, sex, and trait. Hardening consistently improved survival and helped protect fertility under heat stress, including stored sperm fertility in females, a trait where the effect of hardening had previously not been explored. *Spiroplasma* increased survival and male fertility under heat stress, and interacted with plasticity in its native host, suggesting endosymbionts can influence both host upper thermal limits and plastic responses. However, in a novel host, positive effects of *Spiroplasma* and interaction with hardening were limited to male fertility. Nonetheless, *Spiroplasma* and heat hardening generally increased overall fitness in both sexes of both species, translating into reduced heating risk across *D. birchii’s* predicted distribution under current and future climate change. These results underscore the importance of considering multiple, interacting factors, such as plasticity and endosymbionts when predicting species’ responses to climate change and exploring strategies to manipulate thermal limits in ecologically or economically important species. They also highlight the value of fitness_-_based approaches that go beyond survival to capture the full ecological consequences of thermal stress.

## Acknowledgements

We would like to thank Courtney Brown, Nancy M. Endersby-Harshman, Qiong Yang, Harley Thompson, Liam Ferguson, Jackson Young and Vanessa Kellermann for their technical assistance and advice. We would also like to thank the Australian Research Council for financial support to B.v.H. and P.A.R through their Discovery and Fellowship schemes (FT200100025 and DE230100067).

## Author Contributions

Belinda van Heerwaarden Xinyue Gu and Perran A. Ross conceived the ideas and designed methodology; Sian MacDonald, Lucy Hayward and Belinda van Heerwaarden collected the data; Sian MacDonald, Lucy Hayward, Belinda van Heerwaarden and Xinyue Gu analysed the data; Sian MacDonald, Lucy Hayward and Belinda van Heerwaarden led the writing of the manuscript. All authors contributed critically to the drafts and gave final approval for publication.

## Supplementary material

**Table S1.**
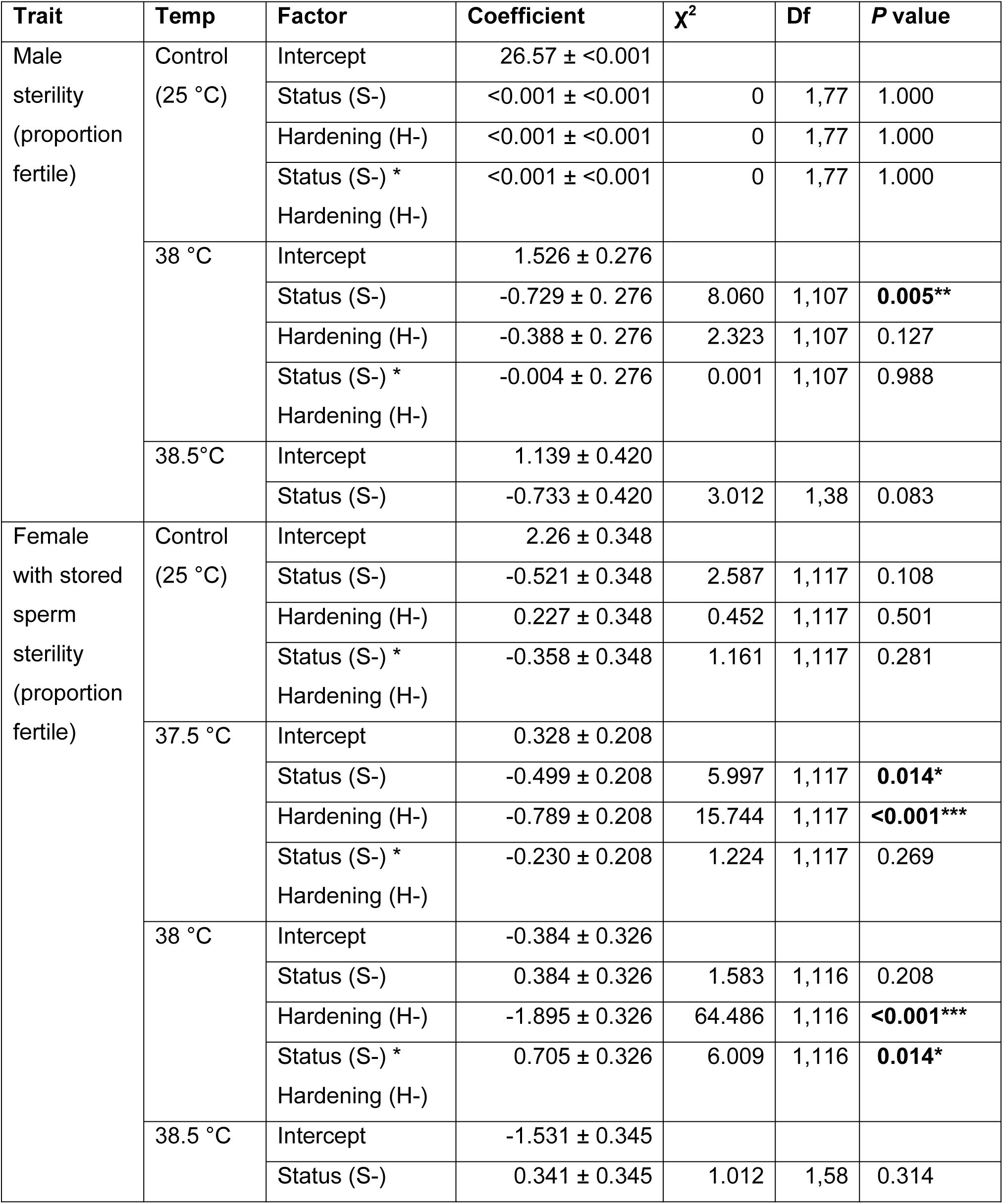
General linear model outputs for *D. hydei* male and female stored sperm sterility after a 1-hour heat shock at different temperatures. H- = unhardened, S- = *Spiroplasma* negative. Binomial distribution, logit link function. When there were issues of separation in the data (i.e. only 0s or 1s in a treatment), we applied Firth’s penalized maximum likelihood estimation using the brglmFit method in R (brglm2 package). * P < 0.05; ** P < 0.01; *** P < 0.001

| Trait | Temp | Factor | Coefficient | $\chi^2$ | Df | P value |
| --- | --- | --- | --- | --- | --- | --- |
| Male sterility (proportion fertile) | Control (25 °C) | Intercept | 26.57 ± <0.001 |  |  |  |
|  |  | Status (S-) | <0.001 ± <0.001 | 0 | 1,77 | 1.000 |
|  |  | Hardening (H-) | <0.001 ± <0.001 | 0 | 1,77 | 1.000 |
|  |  | Status (S-) *<br>Hardening (H-) | <0.001 ± <0.001 | 0 | 1,77 | 1.000 |
|  | 38 °C | Intercept | 1.526 ± 0.276 |  |  |  |
|  |  | Status (S-) | -0.729 ± 0.276 | 8.060 | 1,107 | **0.005\*\*** |
|  |  | Hardening (H-) | -0.388 ± 0.276 | 2.323 | 1,107 | 0.127 |
|  |  | Status (S-) *<br>Hardening (H-) | -0.004 ± 0.276 | 0.001 | 1,107 | 0.988 |
|  | 38.5°C | Intercept | 1.139 ± 0.420 |  |  |  |
|  |  | Status (S-) | -0.733 ± 0.420 | 3.012 | 1,38 | 0.083 |
| Female with stored sperm sterility (proportion fertile) | Control (25 °C) | Intercept | 2.26 ± 0.348 |  |  |  |
|  |  | Status (S-) | -0.521 ± 0.348 | 2.587 | 1,117 | 0.108 |
|  |  | Hardening (H-) | 0.227 ± 0.348 | 0.452 | 1,117 | 0.501 |
|  |  | Status (S-) *<br>Hardening (H-) | -0.358 ± 0.348 | 1.161 | 1,117 | 0.281 |
|  | 37.5 °C | Intercept | 0.328 ± 0.208 |  |  |  |
|  |  | Status (S-) | -0.499 ± 0.208 | 5.997 | 1,117 | **0.014\*** |
|  |  | Hardening (H-) | -0.789 ± 0.208 | 15.744 | 1,117 | **<0.001\*\*\*** |
|  |  | Status (S-) *<br>Hardening (H-) | -0.230 ± 0.208 | 1.224 | 1,117 | 0.269 |
|  | 38 °C | Intercept | -0.384 ± 0.326 |  |  |  |
|  |  | Status (S-) | 0.384 ± 0.326 | 1.583 | 1,116 | 0.208 |
|  |  | Hardening (H-) | -1.895 ± 0.326 | 64.486 | 1,116 | **<0.001\*\*\*** |
|  |  | Status (S-) *<br>Hardening (H-) | 0.705 ± 0.326 | 6.009 | 1,116 | **0.014\*** |
|  | 38.5 °C | Intercept | -1.531 ± 0.345 |  |  |  |
|  |  | Status (S-) | 0.341 ± 0.345 | 1.012 | 1,58 | 0.314 |

**Table S2.**
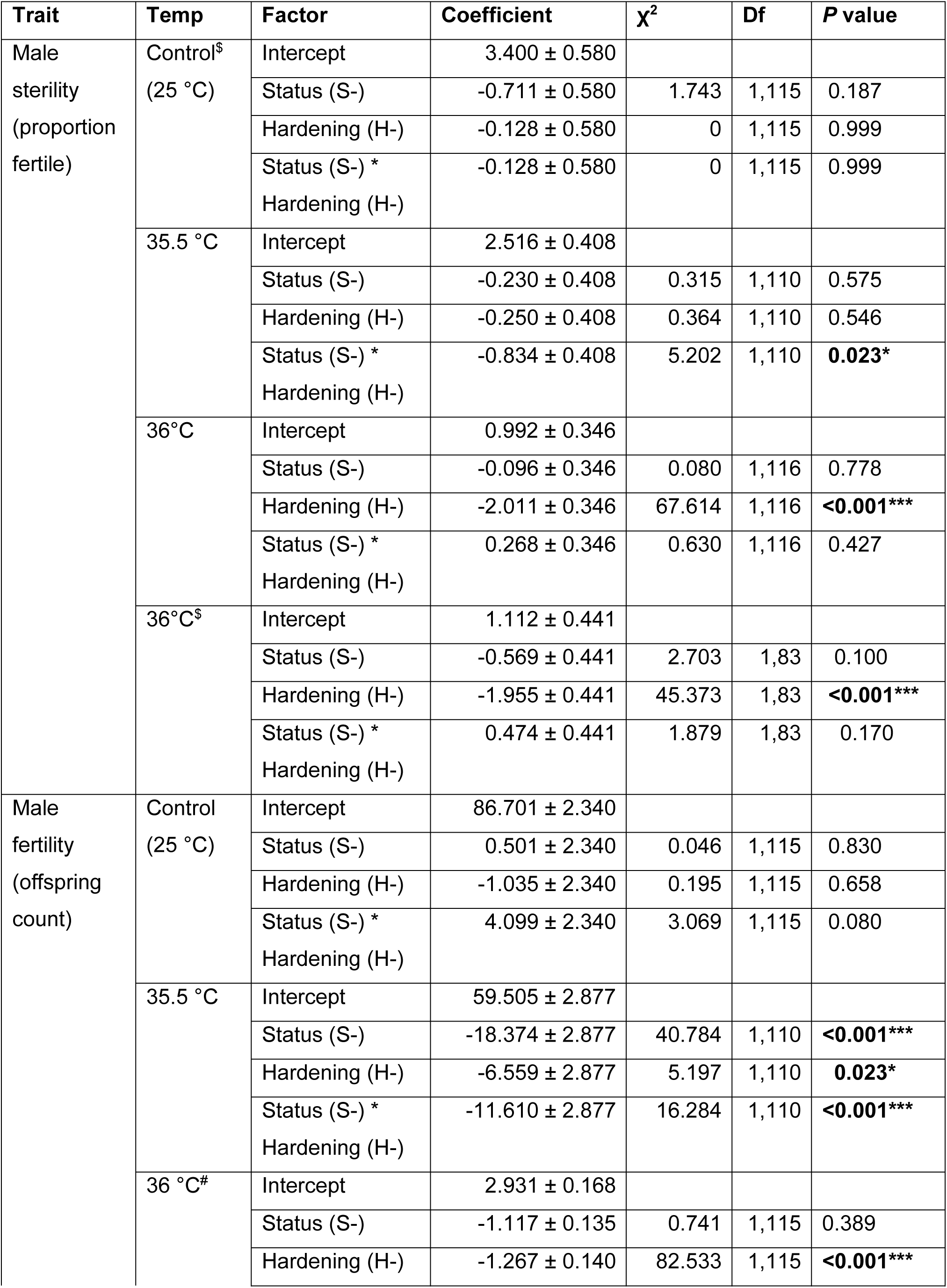
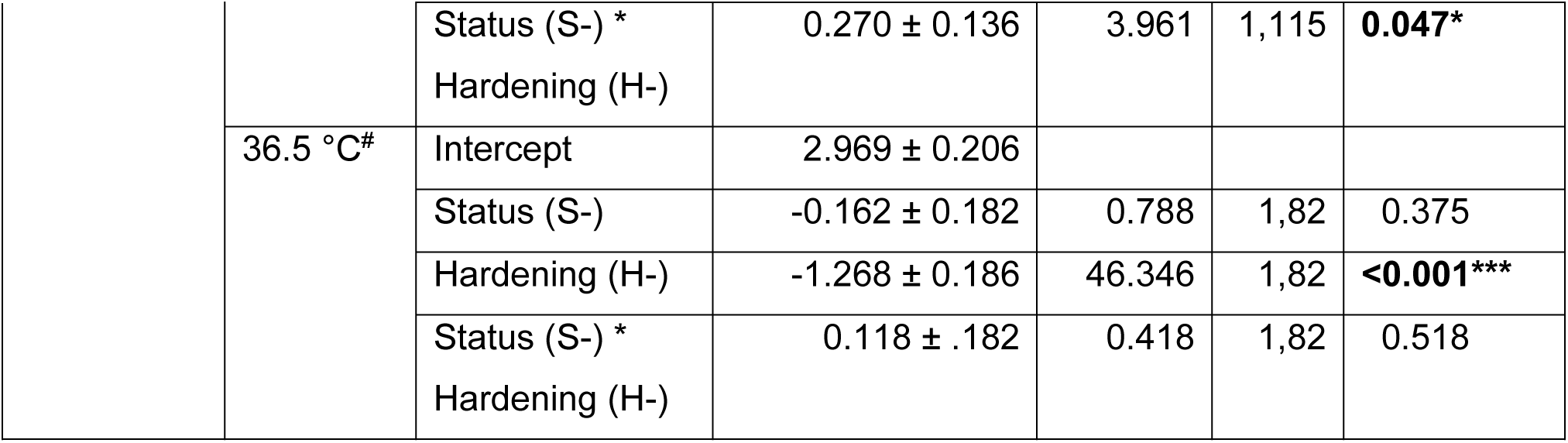
General linear model outputs for *D. birchii* male sterility and fertility after a 1-hour heat shock at different temperatures. H- = unhardened, S- = *Spiroplasma* negative. Binomial distribution, logit link function was used for proportion fertile; gaussian distribution and negative binomial distribution^#^, was used for fertility. When there were issues of separation in the data (i.e. only 0s or 1s in a treatment)^$^, we applied Firth’s penalized maximum likelihood estimation using the brglmFit method in R (brglm2 package).

**Table S3.** General linear model outputs for *D. birchii* female sterility and stored sperm fertility after a 1_-_hour heat shock at different temperatures. H- = unhardened, S- = *Spiroplasma* negative. Binomial distribution, logit link function was used for proportion fertile; negative binomial distribution, was used for stored sperm fertility after heat shock#; and gaussian distribution for remated fertility. When there were issues of separation in the data (i.e. only 0s or 1s in a treatment), we applied firth’s penalized maximum likelihood estimation using the brglmFit method in R (brglm2 package). * P < 0.05; ** P < 0.01; *** P < 0.001

| Trait | Temp | Factor | Coefficient | $\chi^2$ | Df | P value |
| --- | --- | --- | --- | --- | --- | --- |
| Female stored sperm sterility (proportion fertile) | Control (25 °C) | Intercept | 1.386 ± 0.242 |  |  |  |
|  |  | Status (S-) | 0.308 ± 0.242 | 1.655 | 1,116 | 0.198 |
|  |  | Hardening (H-) | 0.518 ± 0.242 | 4.893 | 1,116 | **0.027\*** |
|  |  | Status (S-) *<br>Hardening (H-) | -0.014 ± 0.242 | 0.003 | 1,116 | 0.955 |
|  | 35.5 °C | Intercept | -0.599 ± 0.201 |  |  |  |
|  |  | Status (S-) | 0.160 ± 0.201 | 0.641 | 1,116 | 0.424 |
|  |  | Hardening (H-) | -0.599 ± 0.201 | 9.413 | 1,116 | **0.002\*\*** |
|  |  | Status (S-) *<br>Hardening (H-) | 0.027 ± 0.201 | 0.018 | 1,116 | 0.893 |
|  | 36 °C | Intercept | -1.551 ± 0.249 |  |  |  |
|  |  | Status (S-) | 0.241 ± 0.249 | 0.956 | 1,116 | 0.328 |
|  |  | Hardening (H-) | -0.053 ± 0.249 | 0.046 | 1,116 | 0.830 |
|  |  | Status (S-) *<br>Hardening (H-) | 0.352 ± 0.249 | 2.086 | 1,116 | 0.149 |
|  | 36.5 °C | Intercept | -2.418 ± 0.476 |  |  |  |
|  |  | Status (S-) | 0.221 ± 0.476 | 0.220 | 1,58 | 0.639 |
| Female stored sperm fertility (offspring count) | Control (25 °C) | Intercept | 17.803 ± 1.444 |  |  |  |
|  |  | Status (S-) | 0.892 ± 1.444 | 0.381 | 1,116 | 0.537 |
|  |  | Hardening (H-) | 4.108 ± 1.444 | 8.095 | 1,116 | **0.004\*\*** |
|  |  | Status (S-) *<br>Hardening (H-) | -0.275 ± 1.444 | 0.036 | 1,116 | 0.849 |
|  | 35.5 °C<sup>#</sup> | Intercept | 0.827 ± 0.225 |  |  |  |
|  |  | Status (S-) | -0.131 ± 0.225 | 0.342 | 1,115 | 0.558 |
|  |  | Hardening (H-) | -0.880 ± 0.225 | 15.342 | 1,115 | **<0.001\*\*\*** |
|  |  | Status (S-) *<br>Hardening (H-) | 0.079 ± 0.225 | 0.123 | 1,115 | 0.726 |
|  | 36 °C<sup>#</sup> | Intercept | -0.248 ± 0.326 |  |  |  |
|  |  | Status (S-) | 0.509 ± 0.326 | 2.444 | 1,115 | 0.118 |
|  |  | Hardening (H-) | 0.249 ± 0.326 | 0.585 | 1,115 | 0.444 |
| | | Hardening (H-)<br>Status (S-) * | $-0.198 \pm 0.326$ | 0.369 | 1,115 | 0.544 |
| | 36.5°C<sup>#</sup> | Intercept | $0.039 \pm 0.882$ | | | |
| | | Status (S-) | $0.225 \pm 0.456$ | 0.243 | 1,57 | 0.622 |
| Female<br>remated<br>fertility<br>(offspring<br>count) | Control<br>(25 °C) | Intercept | $63.850 \pm 3.745$ | | | |
| | | Status (S-) | $-1.025 \pm 3.745$ | 0.075 | 1,76 | 0.784 |
| | | Hardening (H-) | $2.175 \pm 3.745$ | 0.337 | 1,76 | 0.561 |
| | | Status (S-) *<br>Hardening (H-) | $0.500 \pm 3.745$ | 0.018 | 1,76 | 0.894 |
| | 35.5 °C | Intercept | $59.912 \pm 4.057$ | | | |
| | | Status (S-) | $-3.637 \pm 4.057$ | 0.804 | 1,76 | 0.370 |
| | | Hardening (H-) | $-1.687 \pm 4.057$ | 0.173 | 1,76 | 0.678 |
| | | Status (S-) *<br>Hardening (H-) | $-2.138 \pm 4.057$ | 0.278 | 1,76 | 0.598 |
| | 36°C | Intercept | $64.438 \pm 3.099$ | | | |
| | | Status (S-) | $-4.963 \pm 3.099$ | 2.538 | 1,76 | 0.111 |
| | | Hardening (H-) | $-6.688 \pm 3.099$ | 4.656 | 1,76 | **0.031\*** |
| | | Status (S-) *<br>Hardening (H-) | $-2.513 \pm 3.099$ | 0.657 | 1,76 | 0.418 |
| | 36.5°C | Intercept | $55.05 \pm 5.03$ | | | |
| | | Status (S-) | $-4.40 \pm 5.03$ | 0.765 | 1,38 | 0.382 |

**Table S4.** General linear model outputs for *D. birchii* male and female fertility in fertile individuals only after a 1-hour heat shock at different temperatures. H- = unhardened, S- = *Spiroplasma* negative. Gaussian distribution was used for males, negative binomial distribution was used for females.

| Trait | Temp | Factor | Coefficient | $\chi^2$ | Df | P value |
| --- | --- | --- | --- | --- | --- | --- |
| Male fertility offspring count (only fertile males) | 35.5 °C | Intercept | 64.580 ± 2.665 |  |  |  |
|  |  | Status (S-) | -18.982 ± 2.665 | 50.750 | 1,99 | **<0.001\*\*\*** |
|  |  | Hardening (H-) | -6.708 ± 2.665 | 6.338 | 1,99 | **0.012\*** |
|  |  | Status (S-) *<br>Hardening (H-) | -9.039 ± 2.665 | 11.508 | 1,99 | **<0.001\*\*\*** |
|  | 36°C | Intercept | 61.537 ± 3.560 |  |  |  |
|  |  | Status (S-) | -12.642 ± 3.560 | 12.614 | 1,69 | **<0.001\*\*\*** |
|  |  | Hardening (H-) | -2.767 ± 3.560 | 0.604 | 1,69 | 0.437 |
|  |  | Status (S-) *<br>Hardening (H-) | 6.984 ± 3.560 | 3.849 | 1,69 | **0.049\*** |
|  | 36.5°C | Intercept | 66.312 ± 5.725 |  |  |  |
|  |  | Status (S-) | 8.996 ± 5.725 | 2.469 | 1,60 | 0.116 |
|  |  | Hardening (H-) | -3.312 ± 5.725 | 0.335 | 1,60 | 0.563 |
|  |  | Status (S-) *<br>Hardening (H-) | 13.004 ± 0.476 | 5.160 | 1,60 | **0.023\*** |
| Female stored sperm fertility offspring count (only fertile females) | 35.5 °C<sup>#</sup> | Intercept | 1.908 ± 0.160 | 141.64 |  |  |
|  |  | Status (S-) | -0.237 ± 0.160 | 2.181 | 1,39 | 0.140 |
|  |  | Hardening (H-) | -0.494 ± 0.160 | 9.523 | 1,39 | **0.002\*\*** |
|  |  | Status (S-) *<br>Hardening (H-) | 0.040 ± 0.160 | 0.063 | 1,39 | 0.802 |
|  | 36 °C<sup>#</sup> | Intercept | -1.508 ± 0.182 | 68.846 |  |  |
|  |  | Status (S-) | 0.309 ± 0.182 | 2.897 | 1,17 | 0.089 |
|  |  | Hardening (H-) | 0.305 ± 0.182 | 2.813 | 1,17 | 0.093 |
|  |  | Hardening (H-)<br>Status (S-) * | -0.486 ± 0.182 | 7.223 | 1,17 | **0.007\*\*** |
|  | 36.5°C<sup>#</sup> | Intercept | 1.709 ± 0.619 | 7.627 |  |  |
|  |  | Status (S-) | 1.303 ± 0.619 | 4.437 | 1,2 | **0.035\*** |

**Figure S1.**
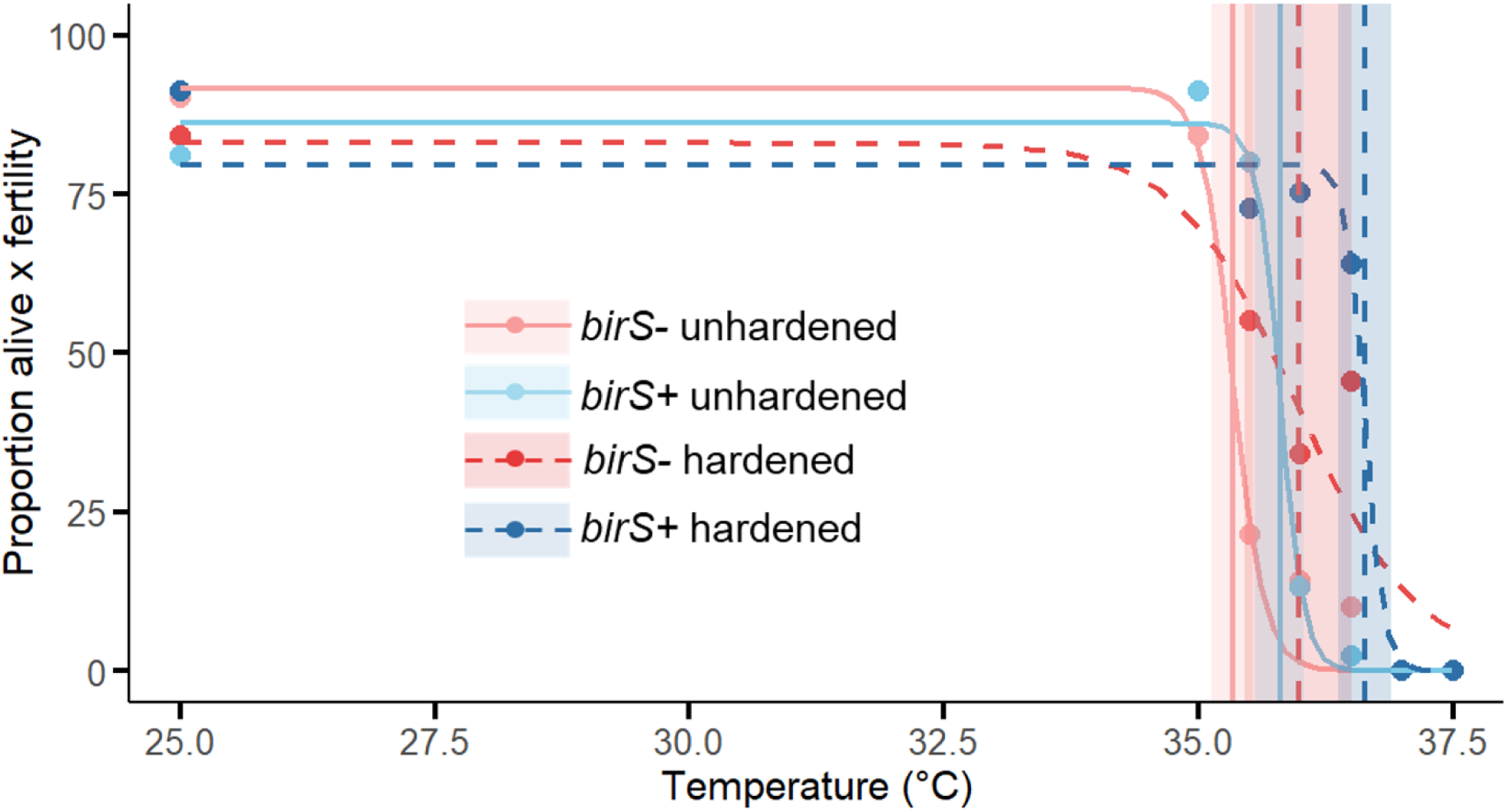
The dose response model used to calculate the temperature at which fitness declines to 50% of controls (25 °C) in adult *D. birchii* males heat hardened 1 hour at 35 °C, with and without *Spiroplasma.* Circles are the average combined survival and fertility measures at each temperate (from Figure 3E) and the fitted lines represent the fitted dose response model. Vertical lines represent are the point at which fitness is 50% of values at 25 °C for each treatment and the shaded lines are the 95% confidence intervals (CIs). Refer to key for *Spiroplasma* and hardening treatments.

**Figure S2.**
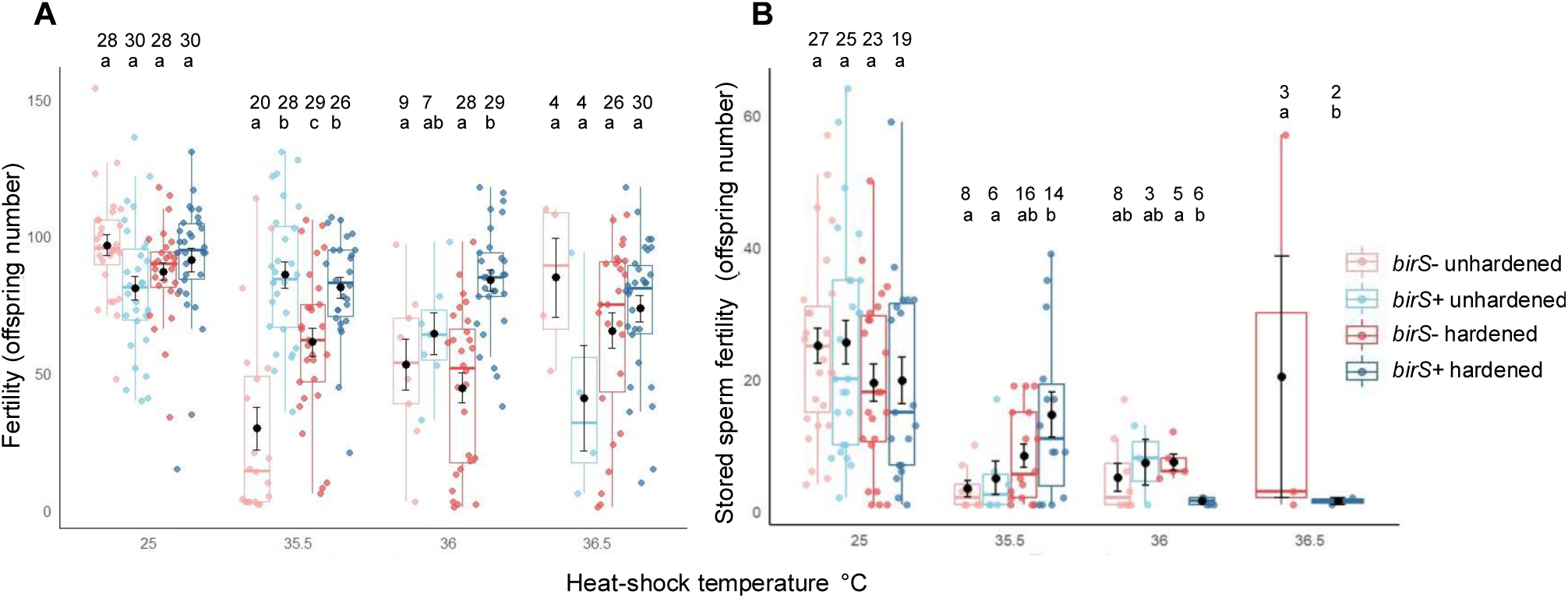
The effect of heat hardening (1 hour at 35 °C) and *Spiroplasma* on *D. birchii* fertility (total offspring count of fertile males/females only) on adult flies exposed to 1-hour heat shocks between 35.5 and 37 °C. Panel A shows effects on males and panel B on mated females. Boxes represent the interquartile range (IQR; 25th to 75th percentile), with the horizontal line indicating the median, whiskers extend to the smallest and largest values within 1.5 × IQR from the lower and upper quartiles, respectively. Jitter points are individual data points. Black dots represent group means, and error bars indicate ±1 standard error of the mean. Letters above indicate treatments that are significantly different from each other (*P* < 0.05, Turkey adjusted). Refer to key for *Spiroplasma* and hardening treatments.

**Figure S3.**
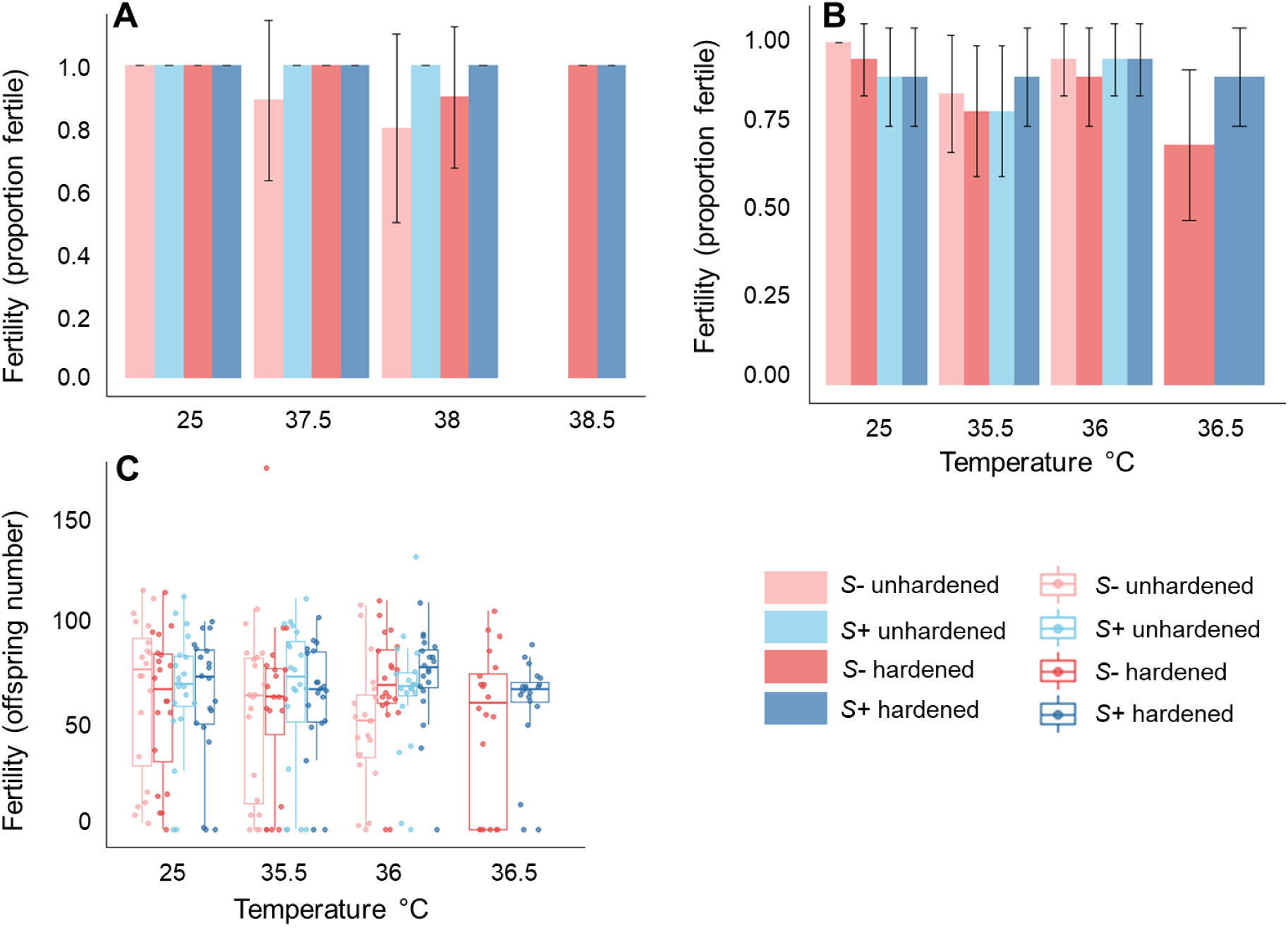
The effect of heat hardening and *Spiroplasma* on fertility of remated heat shocked *D. hydei* and *D. birchii* females. (A) fertility (proportion fertile) of heat shocked *D. hydei* females remated after X days; (B) fertility (proportion fertile) of heat shocked *D. birchii* females remated after X days; and (C) fertility (total offspring number) of heat shocked *D. birchii* females remated after X days. In A and B, bars represent average values, and the capped lines are 95% CIs. In C, the box represents the interquartile range (IQR; 25th to 75th percentile), with the horizontal line indicating the median, whiskers extend to the smallest and largest values within 1.5 × IQR from the lower and upper quartiles, respectively. Jitter points are individual data points. Letters above indicate treatments that are significantly different from each other (*P*<0.05, Turkey adjusted). Refer to key for *Spiroplasma* and hardening treatments.

**Figure S4.**
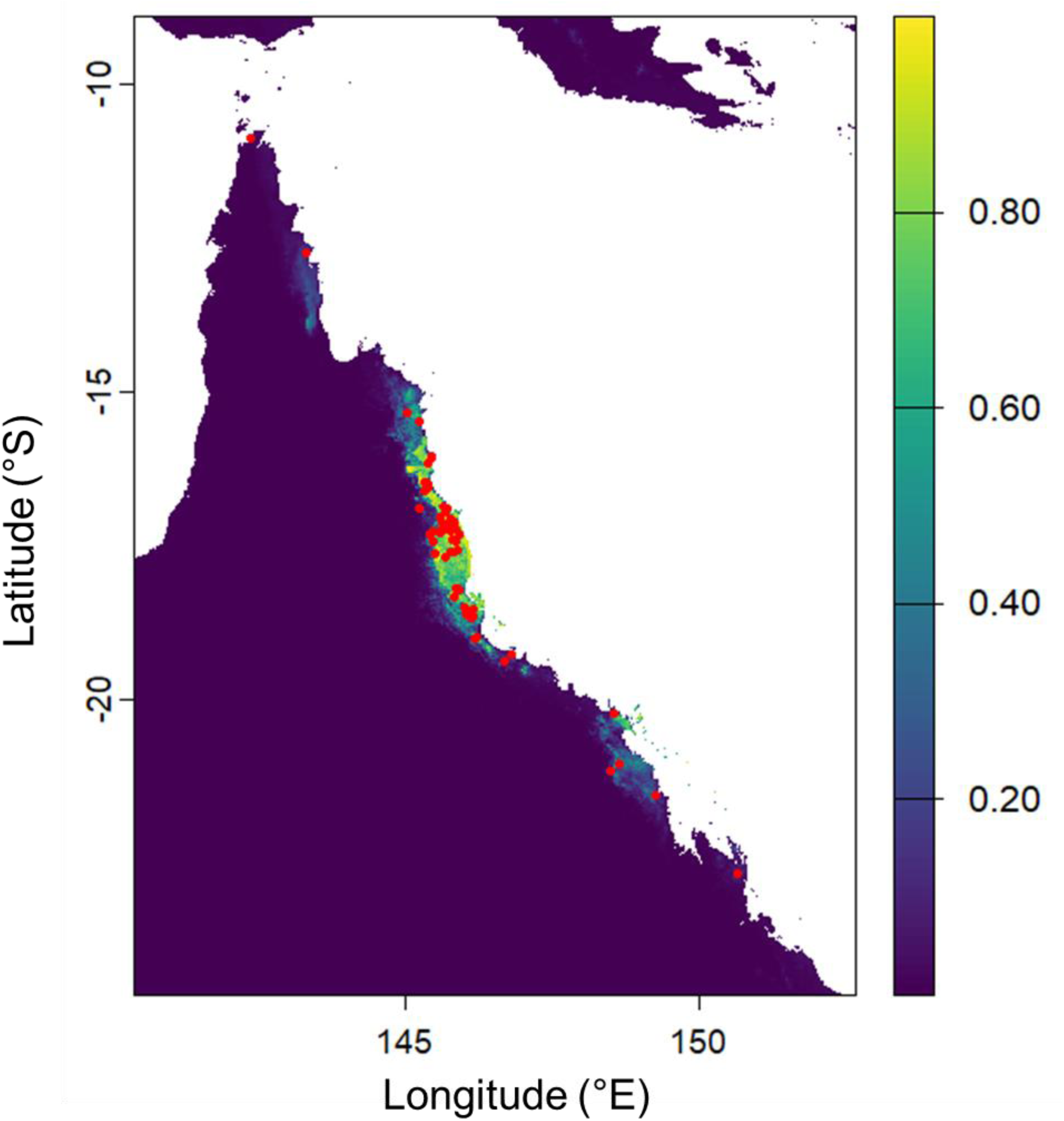
Predicted suitable habitat from MaxEnt for *D. birchii* based on published collection locations in Australia (red dots, Schiffer and McEvey 2006). Predicted habitat suitability values range from 0 (unsuitable) to 1 (highly suitable), with warmer colours indicating greater suitability (see key). AUC = 0.97, 95% CI: 0.959 - 0.987.

**Figure S5.**
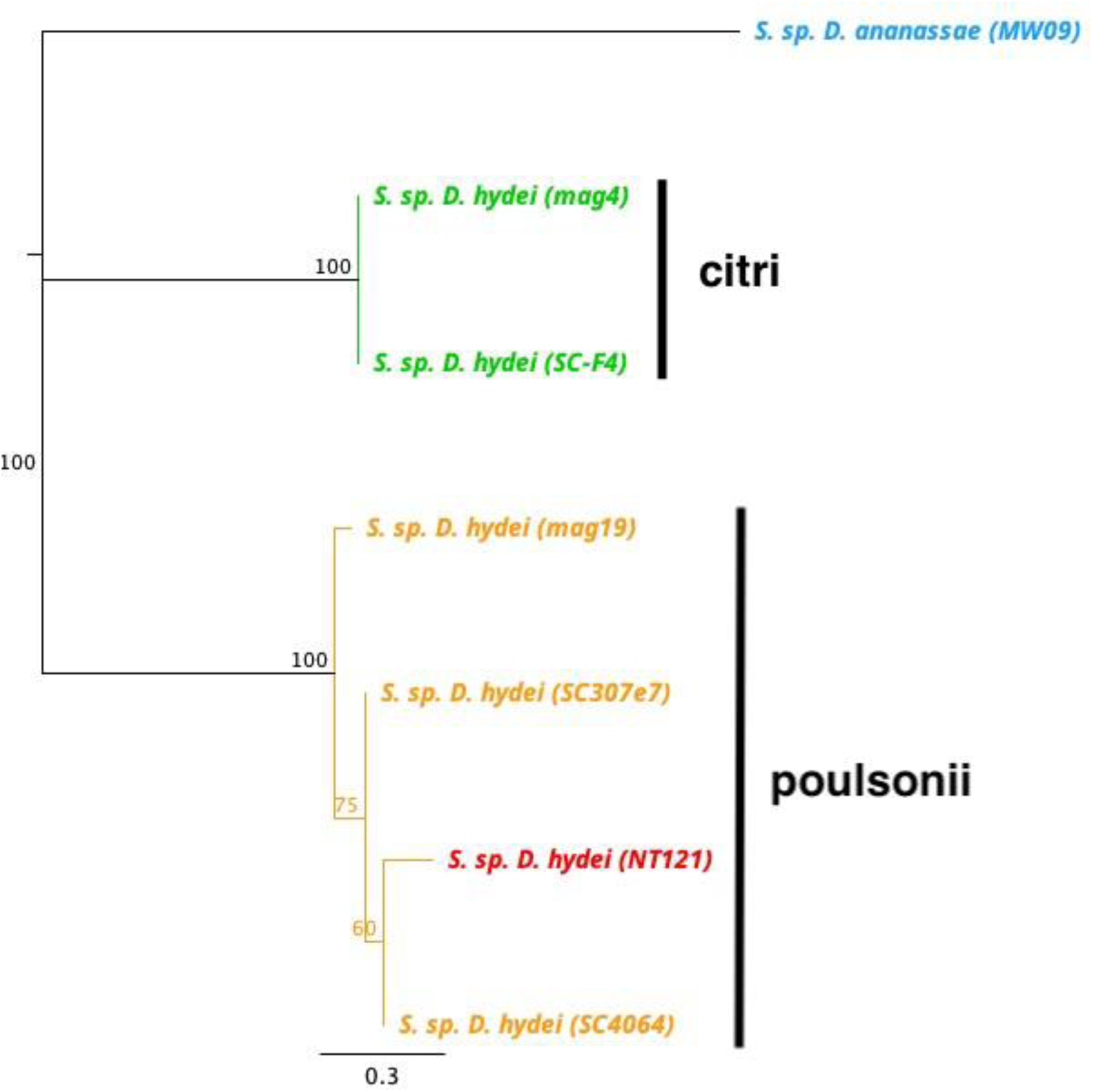
Neighbour-joining phylogeny based on a partial sequence of the 16s rRNA gene from endosymbiont *Spiroplasma*. Different *Spiroplasma* strains are indicated by different colours, where green represents the citri strain and yellow represents the poulsonii strain, excluding the red text which is used to indicate the strain which was sequenced in this study. Sequences from different species of the *Spiroplasma* haplotypes were selected from National Centre for Biotechnology Information (NCBI) for comparison, with *Drosophila ananassae Spiroplasma* strain included as an outgroup.

**Figure S6.**
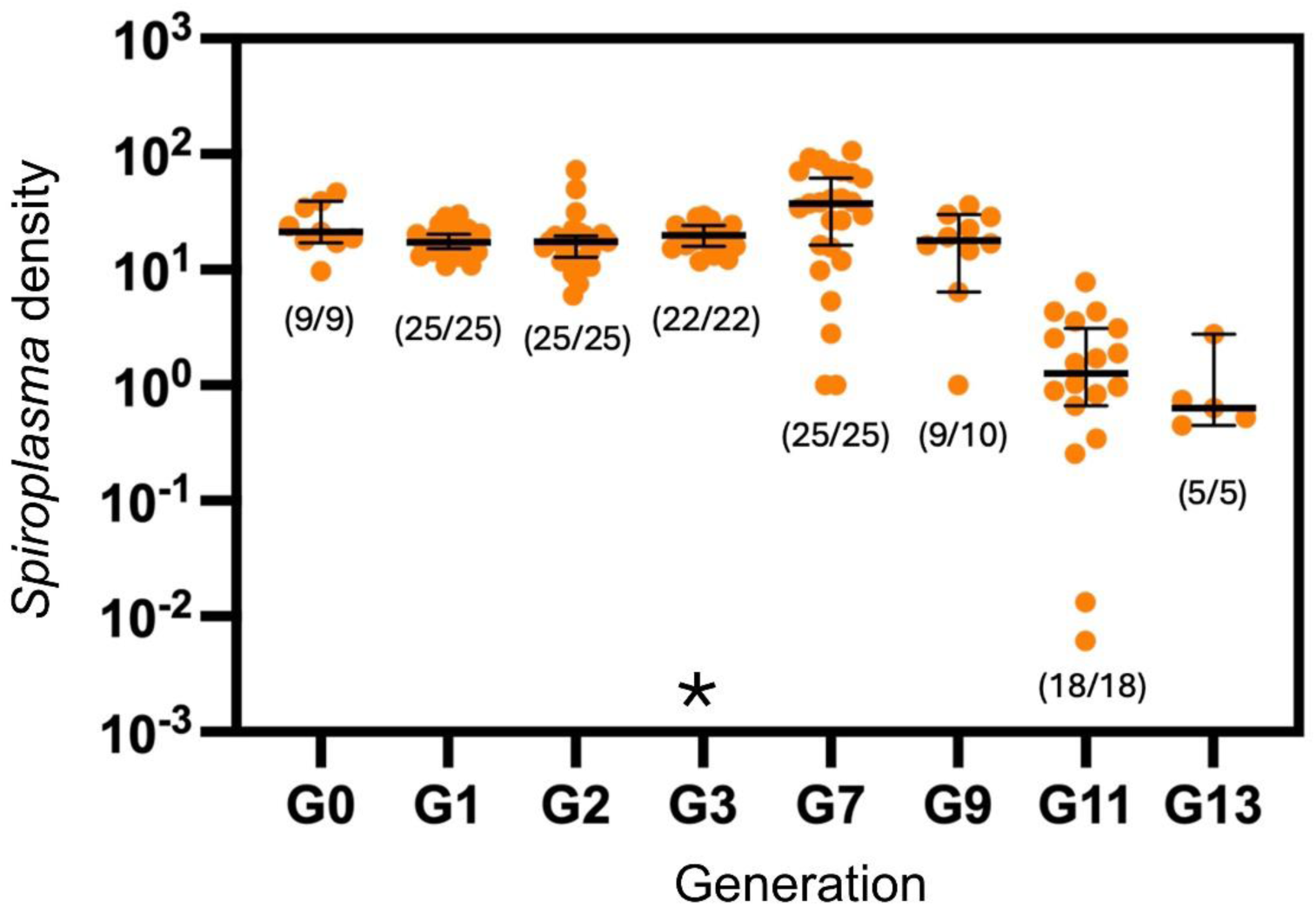
Density of *Spiroplasma* across generations in *Drosophila birchii* that were transinfected with *Spiroplasma* through microinjection. Dots represent data for individual flies, the middle bar shows the median and the error bars show the 95% confidence intervals. Numbers in parentheses represent the *Spiroplasma* frequency for each generation. Generation when matriline selection was stopped is marked by an asterisk.

## Supplementary methods

### *Drosophila* stocks and husbandry

*Drosophila hydei* flies were collected around fruit compost sites at Mountford Farm in Longford, Tasmania in March 2021. Individual field mated females were used to establish iso-female lines (Hoffmann and Parsons 1988). Iso-female lines were maintained in discrete generations and were held in plastic vials, containing 12mL of cornmeal/dextrose media (Hoffmann et al. 1986). These vials were kept in a 25°C temperature-controlled room with a 12-h light: 12-h dark cycle.

*Drosophila birchii* iso-female lines were established from field mated females collected from Kirrama, Queensland from banana baits in March 2021. After 2 generations, a mass-bred population was created by combining 10 males and 10 females from 30 iso-female lines. The massbred population was maintained in discrete generations in 3 x 250 mL bottles (containing 30mL of cornmeal/dextrose media), which were mixed each generation and kept in a 25°C temperature-controlled room with a 12-h light: 12-h dark cycle.

### Screening *D. hydei* lines for *Spiroplasma* presence and strain identification

Chelex DNA extraction and quantitative PCR assays (using methods described in Lee et al. (2012)) were utilised to detect *Spiroplasma* in the *D. hydei* iso-female lines using *Spiroplasma* dnaA primers (dnaA109F 5′-TTAAGAGCAGTTTCAAAATCGGG-3′) and dnaA24 R 5′-TGAAAAAAACAAACAAATTGTTATTACTTC-3′) (Osaka et al. 2013) and *Drosophila* universal primer RpL40 as reference gene (Richardson et al. 2019), and one positive iso-female line (*HyS*+) was chosen for experiments. Relative *Spiroplasma* densities were determined by subtracting the Cp value of the dnaA marker from the Cp value of the RPL40 marker. Differences in Cp were averaged across two to three consistent replicate runs, then transformed by 2n.

To identify the strain of *Spiroplasma* present in the positive *D. hydei* iso-female line, DNA was extracted using a Roche DNA extraction kit according to the manufacturer’s instructions and PCR was used to amplify a section of 16S ribosomal RNA using 23F 5′- CTCAGGATGAACGCTGGCGGCAT-3′) and 1 STR 5′-GGTGTGTACAAGACCCGAGAA-3′) primers (Haselkorn et al. 2009). Amplicons (600bp) were prepared and sent to Macrogen Inc., South Korea, for purification and standard sequencing. Sequences (up to 450bp) were aligned and analysed using Geneious® Prime 2022.2.2 (Biomatters Ltd.) and mapped against reference sequences (GenBank: KC152046.1). Neighbour-joining trees were constructed via the Tamura-Nei model. Sequences from different species of the *Spiroplasma* haplotypes were selected from National Centre for Biotechnology Information (NCBI) for comparison, with *Drosophila ananassae Spiroplasma* strain included as an outgroup (GenBank: KC152046.1). When compared with known *Spiroplasma* sequences present in other *Drosophila* species, it was ascertained that the strain was *Spiroplasma poulsonii* (Figure S5).

### Microinjection of Spiroplasma from D. hydei into D. birchii

We introduced *Spiroplasma* from adult *D. hydei* into adult *D. birchii* through hemolymph microinjection. The *D. birchii* mass-bred used had been in the laboratory for 2 years. Thirty newly emerged (>24 hours old) adult *D. birchii* females were used as the recipients. Twenty adult *D. hydei hy*S+ female flies between five and seven days old were used as the donors of *Spiroplasma*. Flies were initially sedated with carbon dioxide, then kept in an ice box to keep the flies sedated. One single donor fly and recipient fly were removed gently with a fine paint brush from their vials and placed onto a clear 150 mm Petri dish filled with crushed ice to keep flies still during injection. To transfer *Spiroplasma*, a fine glass needle was inserted into the abdomen of the donor *D. hydei* female to collect haemolymph and then directly injected into the thorax of the *D. birchii* female (Nakayama et al. 2015) using a MINJ-1000 microinjection system (Tritech Research, Los Angeles, CA, USA) and a dissecting microscope. One single donor fly had sufficient haemolymph to transfer into approximately three recipient flies.

After the injection, the recovered recipient flies were placed into an empty vial to recover for approximately one hour. After one hour, the recipient flies were then placed into a vial with fresh food and stored under the same conditions as detailed above. They were kept in a group for three days and then nine female survivors were separated into individual vials and given two naturally uninfected males to mate. Flies were kept in vials to mate for one week and then provided fresh food to produce offspring. After two weeks, females were collected and stored individually in ethanol for DNA extraction and qPCR screening for *Spiroplasma*, and the males were discarded. All surviving injected females (N = 9) tested positive for *Spiroplasma* (Figure S6) and six females produced offspring, which were all positive for *Spiroplasma*. Female offspring of each positive female were mated to two uninfected males and allowed to produce offspring for a week. Females were then collected and screened for *Spiroplasma*. This selection process was continued for six generations, to ensure *Spiroplasma* frequency and transmission remained at 100% (Figure S6). These six *Spiroplasma* positive lines were then combined to create a genetically diverse infected line (*birS*+).

### Creating *Spiroplasma* cured lines

Cured *D. hydei* line (*hyS*-) and *D. birchii* line (*birS*-) were generated by creating 20 replicate vials of each line and treating half with antibiotics for two generations. Tetracycline (0.20mg/mL) and Erythromycin (0.16mg/mL) were whisked into the cornmeal-dextrose fly medium once the mixture had cooled to 55°C (Xie et al. 2010, Richardson et al. 2019). Vials with 12mL of antibiotic food media were given to the designated treated lines for two generations. Flies were then given 12mL of regular food media (without Tetracycline/Erythromycin) for another two generations, to recover from potential fitness effects (Xie et al. 2010). To confirm the antibiotics had successfully cured the lines of *Spiroplasma*, qPCR of 10-20 flies per line was used to screen *Spiroplasma* as described above.

### Population density controlling for experiments

Prior to experiments, the density of larvae for the previous and focal generation were controlled, as larval crowding can negatively impact *Drosophila* adult lifespan and heat resistance (Sørensen and Loeschcke 2001). Flies around 5-8 (*D. birchii*) and 10-14 (*D. hydei*) days old were given 24 hours to mate and lay on food media at 25°C. Red food dye was added to this food media to improve visibility of the eggs and was dispensed into either plastic spoons or laying cages. Diluted yeast was added to the surface to encourage the flies to lay eggs. Following the 24-hour laying period, the media was observed under a dissecting microscope and divided into sections of roughly 30 (*D. hydei*) or 40 (*D. birchii*) eggs. The sections were distributed into vials containing 12mL of food media, to raise the subsequent generation for experiments. *Spiroplasma* presence/absence in each line was confirmed prior to experiments as described above.

